# Multistable mechanosensitive behavior of cell adhesion driven by actomyosin contractility and elastic properties of force-transmitting linkages

**DOI:** 10.1101/2023.09.04.554585

**Authors:** Ping Liu, Qiuyu Wang, Mingxi Yao, Artem K. Efremov

## Abstract

The ability of cells to sense the mechanical properties of their microenvironment is essential to many physiological processes. The molecular clutch theory has played an important role in explaining many mechanosensitive cell behaviors. However, its current implementations have limited ability to understand how molecular heterogeneity, such as adhesion molecules with different elasticities, regulates the mechanical response of cell adhesion. In this study, we developed a model incorporating the experimentally measured elastic properties of such proteins to investigate their influence on cell adhesion. It was found that the model not only could accurately fit previous experimental measurements of cell traction force and retrograde actin flow, but also predicted multistablility of cell adhesion as well as a feedback loop between the densities of the extracellular matrix proteins and contractile myosin II motors in living cells. The existence of such a feedback loop was successfully confirmed in experiments. Taken together, our study provides a theoretical framework for understanding how the mechanical properties of adaptor proteins, local substrate deformations and myosin II contractility affect cell adhesion across different cell types and physiological conditions.

## INTRODUCTION

The ability to perceive and adapt to the surrounding environment is a hallmark of living organisms. At the cellular level, a prime example of this is the mechanosensitivity of cells, which can detect and respond to a variety of mechanical signals, such as external force, substrate stiffness and topography [1, 2]. Mechanotransduction is crucial for the regulation of various physiological processes, including cell migration, proliferation, differentiation, and apoptosis [3]. Additionally, aberrant mechanosensing has been implicated in a number of pathological conditions, such as fibrosis, cancer, and cardiovascular diseases [4]. Therefore, a better understanding of cell mechanosensing mechanisms is important for gaining deeper insights into both physiological and pathological processes.

Since the discovery of the mechanosensitive behavior of cell adhesion complexes [1], several potential molecular mechanisms have been proposed to explain this phenomenon. First, it has been shown that formins may play an important role in the force-induced growth of focal adhesions (FAs) by activating the polymerization of actin filaments [1, 5] – a hypothesis supported by single-molecule studies [6, 7]. Furthermore, it was found that applied mechanical load promotes talin-vinculin and *α*-catenin-vinculin interactions, enhancing the linking of integrins and cadherins to the actin cytoskeleton, which is important for cell mechanotransduction [8–13]. In addition, LIM domain proteins, such as zyxin, were also found to be recruited to FAs in response to mechanical stress, further reinforcing and stabilizing them [14]. Finally, it has been proposed that ion channels, such as Piezo 1/2 and L-type calcium channels, may contribute to cell mechanosensing by allowing ion flux in response to force application to a cell, activating downstream signalling pathways [15–17]. Although single-molecule force spectroscopy techniques have greatly improved our understanding of the mechanosensing mechanisms of such proteins [18], much less is known about how their force-dependent response is integrated at the mesoscale level, ultimately directing cell behaviour.

To address this complex problem, various theoretical models have been previously proposed. Prominent among them is the molecular clutch theory, which has been able to link underlying molecular mechanisms to mechanosensitive cell adhesion behaviour and has been successfully applied to model such processes as filopodia extension, traction force generation, and cell spreading [19–30]. This theory proposes that cells form dynamic adhesion links, known as molecular clutches, between the actin cytoskeleton and the extracellular matrix (ECM) through interactions between adaptor proteins, transmembrane adhesion receptors, and the actin cytoskeleton [Figure 1(a)]. These links give rise to a traction force that facilitates cell spreading and migration [19, 20, 28–30]. Furthermore, substrate stiffness modulates the tension and lifespan of molecular clutches via their force-dependent dissociation rates, explaining the mechanosensitive behavior of cell adhesion [20, 24–28].

**FIG. 1.**
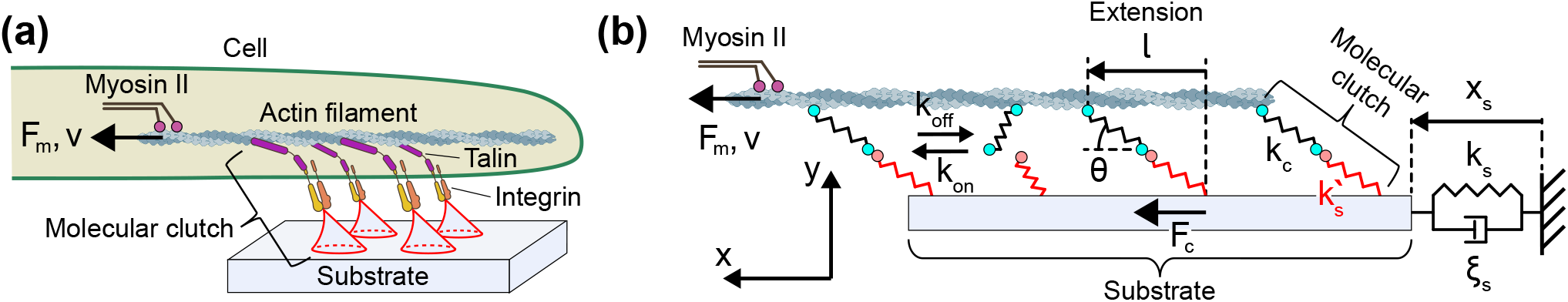
Molecular clutch mechanism. **(a)** Integrins and talins are the main components of adhesion links that form between the actin cytoskeleton and the substrate. Myosin II-generated retrograde flow of actin filaments leads to mechanical stretching of these links, causing local substrate deformations (shown in red). Force-transmitting units, each including an intracellular part (integrin and talin) and an extracellular part (locally deformed substrate), are referred to in this study as ‘molecular clutches’. **(b)** In the model, molecular clutches are represented by composite springs consisting of two parts – a cellular one (*k*_c_) and an extracellular one (*k*_s_^′^). While the latter is assumed to have a linear response to the applied load, the former is considered as a linear spring in the linear clutch model and a nonlinear spring with a talin-like force-response in the talin clutch model. The pulling force (*F*_m_) generated by myosin II motors causes movement of actin filaments at a speed of *v*. This leads to a gradual extension of molecular clutches predominantly along the x-axis parallel to the substrate surface (the contact angle *θ* ≈ 15° [35]), which acquire extension *l*. This, in turn, allows transmission of the myosin II-generated force (*F*_m_) to the substrate, resulting in the cell traction force (*F*_c_), which causes deformation of the substrate (*x*_s_) over a large cell adhesion area. The efficiency of the force transmission is mainly determined by the kinetic behaviour of molecular clutches under mechanical load, which is described by their formation and dissociate rates, *k*_on_ and *k*_off_, respectively. Viscoelastic behaviour of the substrate over a large cell adhesion area is described in the model by a linear spring with the stiffness *k*_s_ and *ξ*_s_ constant, which denotes the viscous properties of the substrate.

Despite its strengths, current implementations of the molecular clutch theory are based on a phenomenological representation of molecular clutches that does not adequately reflect the mechanical and biochemical properties of the underlying molecules. For example, existing models often assume high stiffness of molecular clutches (∼ 1 − 1000 pN/nm [20, 24–27]), resulting in the pre-diction that mechanical stress of molecular clutches is primarily determined by substrate deformations. Yet, such an assumption is inconsistent with single-molecule measurements showing that key elements of molecular clutches, such as talin, are much more elastic due to the presence of long flexible peptide linkers in them, as well as due to the force-induced unfolding of protein domains, being able to extend to more than 200 nm when subjected to a force of several pN [13]. Therefore, although classical model recapitulate mechanosensitive behavior at the cellular level very well, it is difficult to obtain accurate physiological insights at the molecular level with their help, such as estimations of the average tension and force loading rates of molecular clutches and the potential role of the molecular composition and/or mechanical properties in mechanosensitive cell adhesion behaviour, which are the key factor behind cell-type specific mechanical responses of cells.

Furthermore, the ECM is often composed of heterogeneous fibrous materials, such as fibronectin and collagen networks, which are easily deformed at nanometer scales. These types of local deformations have not been taken into account in many previous models, which often treat the substrate as a rigid block on the FAs length scale [20, 24–27]. As a result, such studies often predict large fluctuations in retrograde actin flow [27], which is contrary to the steady retrograde actin flow experimentally observed in filopodia and FAs [20, 26, 31, 32].

These inconsistencies make it difficult to obtain precise molecular insights by comparing experimental data with existing models, hindering understanding of how the mechanical properties of various molecular clutch components, such as talin, kindlin or *α*-actinin, influence the mechanosensitive properties of cell adhesions and how molecular clutch tension dynamics, which have recently been mapped using DNA-based tension sensors [33], relate to cellular response. Moreover, previous theoretical studies have mainly relied on numerical simulations, which has made detailed analysis of the dynamic behavior of cell adhesion very difficult. Addressing these important questions requires an improved theoretical framework based on a more realistic description of the key molecular clutch components.

In this work, we developed a semi-analytical model of cell adhesion based on the molecular clutch theory, incorporating local substrate deformations at adhesion sites of molecular clutches as well as the experimentally measured force-response of the main molecular clutch component, talin. We demonstrate that the developed model is able to fit experimental data on cellular mechanical response equally well or better than previous molecular clutch models, while providing greater physiological insight into the mechanical responses of individual molecular clutches. Model predictions suggest that the elastic properties of molecular clutch components are one of the key factors affecting the mechanical response of living cells, and that molecular clutch dynamics are dominated by steady-state behavior that can exhibit bifurcations in certain scenarios. Furthermore, it is shown that the model can potentially allow to understand more complex scenarios, such as the maturation of FAs, as well as the different mechanical responses of molecular clutches with various component compositions.

## METHODS

### A. Linear molecular clutch model

To address the above issues, we developed a semi-analytical approach based on consideration of the long-term behaviour of molecular clutches, incorporating into the model experimentally measured force-response of adaptor proteins and local substrate deformations at the adhesion sites of molecular clutches. Specifically, we used the model schematically shown in Figure 1(b) in which molecular clutches were represented by springs consisting of two parts: 1) cellular, corresponding to adaptor proteins, and 2) extracellular, describing local substrate deformations at the adhesion sites of molecular clutches. In the simplest case, these two parts can be represented by linear springs with spring constants *k*_c_ and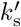, respectively.

In addition to local substrate deformations at the adhesion sites of molecular clutches, the model also took into account the average substrate deformation over a much larger area of cell adhesion of radius *R* (*R* ≈ 1.7 *µ*m [26]), encompassing many molecular clutches, which was described by the spring constant *k*_s_ shown in Figure 1(b).

Using the theory of elasticity, it can be shown that both 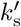 and *k*_s_ are related to the Young’s modulus of the substrate (*E*) as [34]:

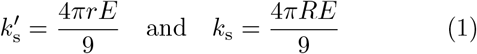

Where *r* is the characteristic radius of the adhesion site of a single molecular clutch.

To describe the dynamic state of the system, all sites available for the molecular clutch formation were divided into two groups in the model: those that are occupied by molecular clutches (’on’ state) and those that are not (’off’ state).

In addition, each molecular clutch was characterized by its own deformation, resulting from to the mechanical load *F*_m_ generated by myosin II motors, see Figure 1(b). Super-resolution studies of the nanoscale architecture of FAs show that the average contact angle of the intracellular part of molecular clutches with the cell membrane / substrate surface, *θ* [Figure 1(b)], is quite small (*θ* ≈ 15° [35]). Thus, molecular clutches experience much greater deformation along the x-axis parallel to the substrate surface compared to the y-axis perpendicular to it. As a result, to a first approximation, the physical state of each molecular clutch can be described by its extension, *l*, along the x-axis parallel to the substrate surface, see Figure 1(b). Accordingly, the state of the entire system can be represented by a time-dependent probability distribution, *p*_on_(*l, t*), to observe a molecular clutch with the extension *l* at time *t* at each site available for the molecular clutch formation. This distribution satisfies the following normalization condition:

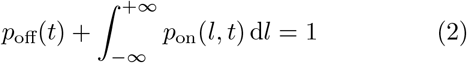

Where *p*_off_(*t*) is the time-dependent probability of the off-state. Molecular clutch extension can be positive or negative depending on the direction of the molecular clutch along the x-axis.

In the model, the rate of molecular clutch formation was described by the first-order rate constant 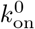. Furthermore, it was assumed that, immediately after assembly, molecular clutches are in a mechanically relaxed state with zero extension along the x-axis (*l* = 0). As a result, the extension-dependent assembly rate of molecular clutches, *k*_on_(*l*), was expressed in terms of the Dirac *δ*-function, *δ*(*l*), as:

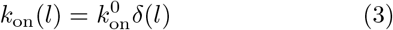

Where 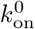 is a constant.

Retrograde flow of actin filaments created by myosin II motors causes gradual stretching of molecular clutches. This, in turn, leads to an increase in their deformation, causing subsequent dissociation. It was assumed in this study that dissociation of molecular clutches is mainly driven by detachment of integrins from their ECM ligands, such as fibronectin. Indeed, previous studies suggest that this may be the weakest point of molecular clutches, as bonds between other molecular clutch components appear to be greatly strengthened by applied mechanical load [36, 37]. Since integrin-ECM bonds have previously been shown to possess catch-bond behaviour [38], the dissociation rate of molecular clutches (*k*_off_) was approximated by the following two-term exponential formula, which was found to fit experimental data quite well [Figure S1, SI]:

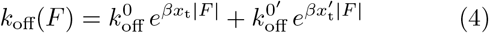

Where *x*_t_ and 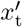 are two transition state distances characterizing the force-dependent behaviour of the integrin-ECM catch-bond, and 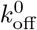 and 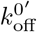 are the corresponding dissociation rates at zero mechanical load. *β* = 1*/k*_B_*T* is the reciprocal of the thermodynamic temperature. |*F*| is the absolute value of the tension of the bond. In the case of molecular clutches represented by two-part linear springs:

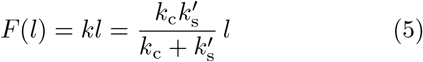

Here 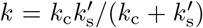 is the resulting spring constant of molecular clutches consisting of the two parts: extracellular 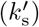 and intracellular (*k*_c_) [Figure 1(b)].

Given the probability distribution *p*_on_(*l, t*), the total tension of molecular clutches, or what is the same, the traction force created by molecular clutches on the substrate, *F*_c_(*t*) [Figure 1(b)], can be found as:

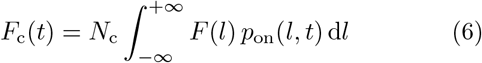

Where *N*_c_ is the total number of sites available for the formation of molecular clutches in a cell adhesion area. Recent experimental studies show that the average density of talin, and therefore likely the density of sites available for molecular clutch formation, is very similar in both small and large FAs [39], suggesting that it is independent of the size of FAs. As a result, it can be concluded that *N*_c_ should be proportional to the cell adhesion area: *N*_c_ = *πR*^2^*σ*_c_, where *σ*_c_ is the average density of sites available for the formation of molecular clutches in the cell adhesion area with a characteristic radius *R*.

According to the Newton’s third law, the total tension of engaged molecular clutches is equal to the force generated by myosin II motor proteins, *F*_m_(*t*) = *F*_c_(*t*). Following previous studies [20, 24–27], the resisting effect of such a load on the movement rate of myosin II motors, *v*(*t*), was approximated by a linear force-velocity relation:

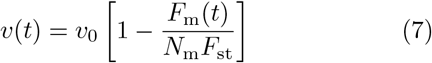

Here *F*_st_ is the stalling force of a single myosin II motor. *v*_0_ is the movement rate of myosin II motors at zero mechanical load. *N*_m_ is the total number of myosin motors pulling actin filaments in a cell adhesion area. Since previous studies suggest that myosin II motors exert a local pulling effect on the actin cytoskeleton in the cell cortex [32], it was assumed in the model that *N*_m_ is proportional to the cell adhesion area: *N*_m_ = *πR*^2^*σ*_m_, where *σ*_m_ is the average surface density of myosin II motors near the cell-substrate interface.

Experimental studies show that actin filaments exhibit retrograde movement at nearly constant rates (*v*(*t*) ≈ const) [20, 26, 31, 32]. Thus, it can be concluded that myosin II motors and the entire ensemble of molecular clutches function close to a steady-state in cells. This makes it possible to derive a system of simple equations that describe the behaviour of the molecular clutch system. Namely, by combining the above formulas, it can be shown that the dynamics of the entire molecular clutch system in the general case is given by Eq. (A10) from Appendix A, SI, which has the following stationary solution that describes the long-term behaviour of the entire ensemble of molecular clutches and mysoin II motor proteins [Appendix A, SI]:

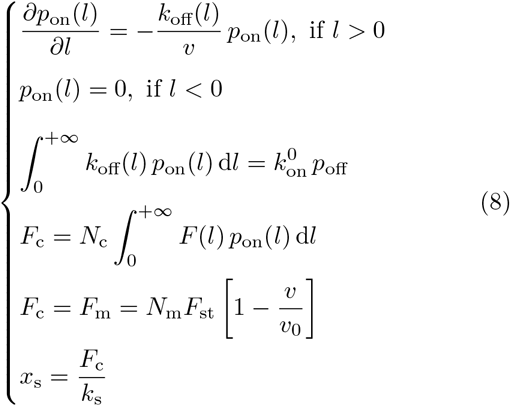

Here *p*_on_(*l*) = ⟨*p*_on_(*l, t*)⟩ _*T*_ is the long time-averaged probability distribution. *k*_off_(*l*) = *k*_off_(*F* (*l*)). *x*_s_ and *k*_s_ are the average mechanical deformation of the substrate and a constant describing the elastic properties of the substrate over the cell adhesion area [Figure 1(b)].

To find out the long-term behaviour of the molecular clutch system, one needs to solve Eq. (8) in terms of *p*_on_, *F*_c_ and *v*. From Eq. (8) it is clear that it can be solved by finding the intersections of *F*_c_(*v*) and *F*_m_(*v*) graphs, which allows not only to construct a simple geometric interpretation of the solutions of Eq. (8), but also to assess their stability, see Figure 2.

**FIG. 2.**
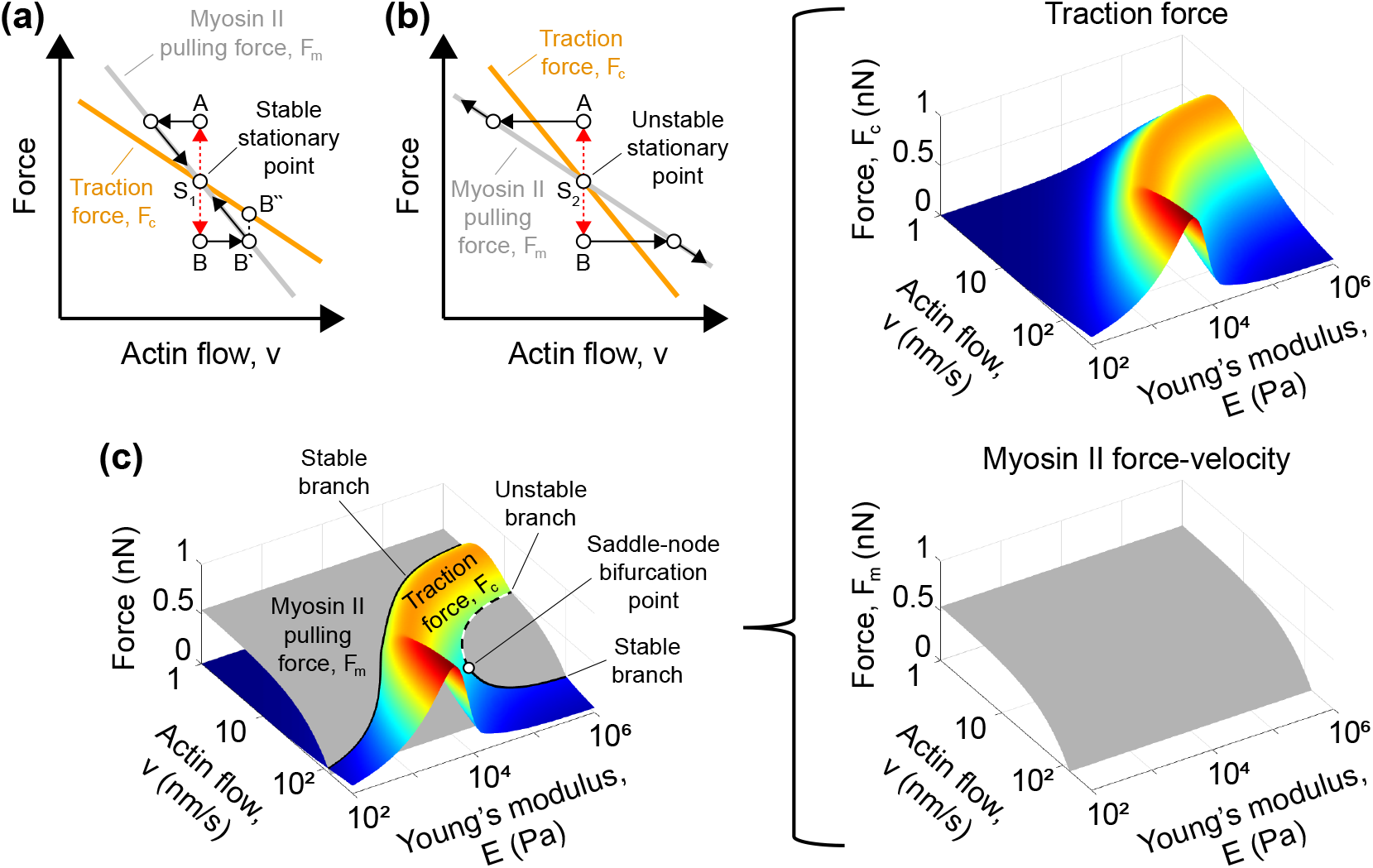
Geometric interpretation of the force balance in the molecular clutch system. **(a, b)** Intersections of the traction force curve, *F*_c_(*v*), and the force-velocity curve of myosin II, *F*_m_(*v*), correspond to the stationary points (*S*_1_ and *S*_2_) described by Eq. (8), at which the molecular clutch system reaches mechanical equilibrium. Stability of trajectories near such points is largely determined by the slopes of the cell traction and myosin II force-velocity curves. Namely, if the latter is greater than the former [panel (a)], trajectories in the neighbourhood of the corresponding stationary point are stable (point *S*_1_). That is, small perturbations do not cause them to move far from the stationary point. On the other hand, if the slope of the myosin II force-velocity curve is smaller than that of the traction force, trajectories near the stationary point will exhibit unstable behaviour [panel (b)]. As an example, consider point *B* in panel (a). The initial deviation of the system away from the stationary point *S*_1_ (for example, due to dissociation of several molecular clutches) will first lead to an increase in the retrograde actin flow, since myosin II motors have to work against a smaller resisting force. However, to maintain a steady flow of actin at a rate corresponding to the new state (point *B*^′^), myosin II motors must, in the long run, balance the resisting traction force created by molecular clutches, which is determined by point *B*^′′^. Since the pulling force generated by myosin II motors is less than the stationary traction force (i.e., point *B*^′^ is below point *B*^′′^), this will cause the retrograde actin flow to slow down, which will eventually bring the system closer to the stationary point *S*_1_. Similar considerations apply to the rest of the points shown in panels (a) and (b), demonstrating the long-term stability of trajectories near the stationary point *S*_1_ and the instability of trajectories near the stationary point *S*_2_. Numerical simulations of the dynamic behaviour of the molecular clutch system show good agreement with the above physical considerations, see Figure 7 and Figure S6(b), SI. **(c)** Example of 3D plots of the cell traction force (*F*_c_, coloured surface) and myosin II-pulling force (*F*_m_, gray surface) as a function of the retrograde actin flow (*v*) and Young’s modulus of the substrate (*E*). Black solid curves formed by the intersection of the two surfaces on the left panel indicate stationary points of the system that attract trajectories to their neighbourhood (stable branches), whereas the black dashed curve indicates unstable stationary points (unstable branch). White dot designates a saddle-node bifurcation point where the stable and unstable branches merge together. All cell traction and retrograde actin flow curves shown in the Results section were obtained by projecting the stable and unstable branches of 3D graphs like those displayed in panel (c) onto the vertical and horizontal planes, respectively.

While *F*_m_(*v*) can be directly calculated from the third line of Eq. (8), computation of *F*_c_(*v*) requires knowledge of *p*_on_(*l*) probability distribution, which can be found from the first three lines of Eq. (8) [Appendix B, SI]:

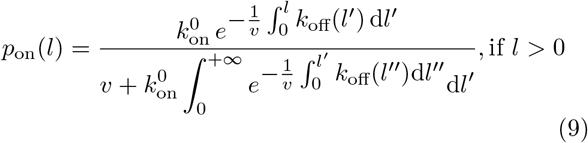

Knowing *p*_on_(*l*) distribution, one can easily obtain the total tension of the engaged molecular clutches (*F*_c_) from Eq. (8) as a function of the retrograde actin flow speed (*v*), and the traction stress (*P*) exerted by a cell to the substrate:

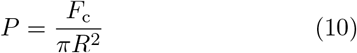

In the special case of slip-bonds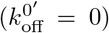, Eq. (9) reduces to a simple analytic formula [Appendix B, SI], which is in good agreement with previous studies [21– 23]:

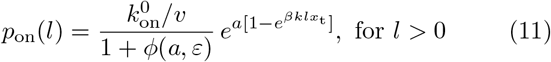

Where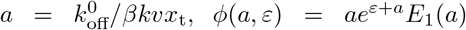 and 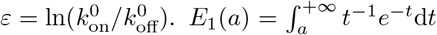 is the exponen-tial integral function.

Substituting Eq. (11) into Eq. (6), we get:

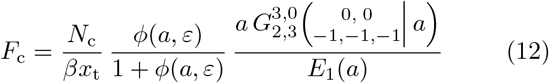

Here 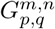 is the Meijer’s G-function, see Figure S3, SI. Finally, it should be noted that although above we considered only molecular clutches represented by linear springs, the resulting Eq. (8)-Eq. (10) and Eq. (A10) are very general in nature, allowing an arbitrary force-response of molecular clutches to be used in the model. For example, to describe plastic deformations of molecular clutches caused by force-induced unfolding of their globular protein domains, it is sufficient to introduce one additional parameter into the model – the yield strength of molecular clutches, changing Eq. (5) to:

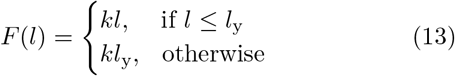

Where *l*_y_ is the molecular clutch extension corresponding to the yield strength.

This coarse-grained approach is most suitable for description of molecular clutches with unknown architecture / protein structure.

On the other hand, if the structure and mechanical stability of the main molecular clutch components are known, more detailed information can be obtained compared to the approach described above by considering all possible transitions between the different folded / unfolded states of molecular clutches. We utilized this method to account for the highly nonlinear behaviour of talin molecules and the force-induced unfolding of the mechanosensitive talin domain (R3) observed in single-molecule experiments [13], see Appendix C, SI.

It is important to note that almost all model parameters can be measured experimentally, see Table T1, SI. As for the remaining parameters, by simultaneously fitting experimental data obtained under different experimental conditions (wild-type or talin 1,2 knockdown cells treated with myosin II inhibitor or grown on substrates coated with different concentrations of fibronectin), the average number of fitting parameters per curve can be reduced to one or two, making the model very robust. A list of key model fitting parameters used in this study can be found in Table 1 in the main text.

**TABLE 1.** Values of fitting parameters in the linear KD, linear WT and talin WT models ^(1)^.

| Parameter | Model <sup>(2)</sup> |  |  | Description |
| --- | --- | --- | --- | --- |
|  | Linear KD | Linear WT | Talin WT |  |
| $k_{\text{on}}^0$ | $0.005 \text{ s}^{-1}$ | $0.23 \text{ s}^{-1}$ | $0.3 \text{ s}^{-1}$ | Assembly rate of molecular clutches in a mechanically relaxed state. The estimated values are in fairly good agreement with the experimentally measured rates of formation of talin-based molecular clutches in *Xenopus laevis* cells ($0.03 - 0.06 \text{ s}^{-1}$ [42]). |
| $\sigma_{\text{m}}$ | $71 \mu\text{m}^{-2}$ | $85 \mu\text{m}^{-2}$ | $82 \mu\text{m}^{-2}$ | Density of myosin II motor proteins near the cell-substrate interface. The estimated values are close to the experimentally measured density of myosin II filaments ($\sim 2 - 3 \mu\text{m}^{-2}$ [32, 46]) multiplied by the average number of individual myosin molecules in each myosin II filament ($\sim 30$ [57]). |
| $k_{\text{c}}$ | $1.6 \text{ pN/nm}$ | $0.05 \text{ pN/nm}$ | | Spring constant of the intracellular part of molecular clutches (linear KD and linear WT models only). Experimentally measured spring constant of talin in the physiologically relevant range of $0 - 10 \text{ pN}$ is $\sim 0.1 \text{ pN/nm}$, see Table T3, SI. |
| $r$ | $26 \text{ nm}$ | $0.8 \text{ nm}$ | $0.6 \text{ nm}$ | Characteristic radius of the adhesion site of a single molecular clutch. |
| $l_{\text{y}}$ | $150 \text{ nm}$ | $> 300 \text{ nm}$ | | Molecular clutch extension corresponding to the yield strength (linear KD and linear WT models only). The value of this model parameter did not have a strong effect on the results in the case of the linear WT model. |

<sup>1</sup> A complete list of model parameters can be found in Table T1, SI.
<sup>2</sup> In the linear KD and linear WT models describing MEF Talin 1 KO, Talin 2 shRNA cells and MEF WT cells, respectively, molecular clutches were represented by two-part linear springs up to the yield point, $l_{\text{y}}$ , see Eq. (13) in the Methods section. In the talin WT model describing MEF WT cells, the intracellular parts of molecular clutches were represented by nonlinear springs with a talin-like force-response, whereas the extracellular part was represented by a linear spring, see Methods and Appendix C, SI.

### B. Cell lines, constructs and experimental data processing

#### Cell culture, constructs and transfection

NIH-3T3 and HeLa cells were obtained from the laboratory of Dr. Qin Peng (Shenzhen Bay Laboratory, China). All cell lines were grown in Dulbecco’s Modified Eagle Medium (DMEM, Gibco, C11995500BT) supplemented with 10% fetal bovine serum (FBS, Gibco, 10099-141C) and 1% penicillin/streptomycin (Gibco, 15070-063) at 37°C, 5% CO_2_. To visualize myosin II motors, the cells were transfected with a plasmid encoding myosin regulatory light chain (RLC)-GFP (gift from Dr. Alexander Bershadsky, Mechanobiology Institute, Singapore) using jetPRIME buffer (Polyplus, 712-60) and jetPRIME reagent (Polyplus, 114-15) according to the manufacturer’s instructions. Following the transfection, cells were seeded onto 35 mm glass bottom dishes coated with 1 *µ*g/ml, 10 *µ*g/ml or 100 *µ*g/ml fibronectin (Solarbio, F8180) dissolved in Dulbecoo’s Phosphate Buffered Saline (DPBS, Gibco, C14190500BT). The next day after transfection, cells were fixed for 15 min with 4% paraformaldehyde solution (Beyotime, P0099), followed by washing with DPBS (3 times), after which cells were imaged.

#### Fluorescence microscopy

Fluorescent specimens were imaged using ultra-high resolution microscope Elyra 7 (ZEISS) equipped with Plan Apochromatic 63 ×/1.4 Oil (DIC) objective, which was controlled by ZEN (black edition) software. The images were captured with pco.edge sCMOS camera (Lattice SIM module) and processed by Lattice SIM2 structured illumination technique (horizontal resolution: 60 nm, axial resolution: 200 nm). Fluorescence excitation was performed using 488 nm, 500 mW OPSL and 561 nm, 500 mW OPSL.

#### Image analysis and statistics

Collected images of RLC-GFP labelled myosin II stacks were analysed with the help of ImageJ-win64 software. To this aim, for each cell, several random regions near the cell corners were selected, and automatic thresholding was applied using built-in ’Auto Threshold’ function of ImageJ in order to map the contours of myosin II stacks. The resulting images were further processed with ’Analyze Particles’ function, which provided the total number and sizes of the detected myosin II stacks. After discarding all particles with an area less then 0.01 *µ*m^2^, the average density and the average size of myosin II stacks, as well as corresponding standard errors of the mean, were calculated. All experimental results presented in the article are representative of three independent experiments. Statistical analysis of the data was carried out using the non-parametric Mann-Whitney test.

## RESULTS

### Semi-analytical model based on the physiological molecular mechanics of cell adhesion complexes

To cope with the limitations of existing models based on the molecular clutch theory, we developed a semi-analytical theoretical framework consistent with the experimentally observed force-response of molecular clutch components at the single-molecule level [12, 13] and experimentally measured tension [39, 40] and loading rates of molecular clutches [33]. This allows for a comprehensive study of the dynamic aspects of cell adhesion me-chanics, providing new insight into the role of different cellular elements in shaping mechanosensitive cell adhesion behaviour.

In the model, individual molecular clutches are represented by two-part springs: one part describes cell adaptor proteins connecting adhesion receptors to the cytoskeleton, and the other part depicts local deformations of the substrate at the attachment sites of the adhesion receptors, see Methods section and Figure 1. The latter part of molecular clutches can be related to the substrate rigidity through a spring constant, the value of which can be obtained from the theory of elasticity [Eq. (1)].

In addition, the model distinguishes between ‘occupied’ (on) and ‘free’ (off) states of sites available for the formation of molecular clutches, and quantifies the deformation of each molecular clutch under the influence of mechanical load created by the contractile activity of myosin II motors. This allows the probability distribution of molecular clutch extension to be described over time, making it possible to calculate the total traction force exerted by molecular clutches on the substrate, which results from the molecular clutch-mediated transmission of the pulling force generated by myosin II motors, see Figure 1(b) and Eq. (6). The efficiency of this force transmission, which is determined by the dynamics of molecular clutch formation and dissociation, is a key element for understanding mechanosensitive cell adhesion behaviour, as can be seen from the master equation describing the time evolution of the molecular clutch system, see Eq. (A10) in Appendix A, SI.

Utilizing a few physiologically relevant fitting parameters (Table 1), our model can fit well with previously published experimental data, such as retrograde actin flow and cell traction measured as a function of the substrate Young’s modulus [26], see the results below. This makes it possible the model to be used to plot cell adhesion stability diagrams and extract detailed information about molecular clutch dynamics at the single-molecule level from experimental data obtained on cells, including the average tension and conformational changes of molecular clutches, providing new insight into the key factors regulating FA behavior.

### Potential effect of talin force-response on cell adhesion

Talin was shown to be a major component of molecular clutches in FAs, and its deletion was found to cause dramatic changes in mechanosensitive cell adhesion behavior, which was previously explained to be due to the force-induced enhancement of FAs by talin-vinculin interactions [26]. However, this view conflicts with multiple experimental studies showing that vinculin is not an essential component of force-transmission pathways mediated by molecular clutches [39–41], suggesting that other factors, such as the talin force-response, may play a role in regulating mechanosensing behavior of molecular clutches.

To this aim, in our model we treated the intra-cellular part of molecular clutches as a nonlinear spring with a force-response corresponding to that of talin measured in single-molecule studies [13], including the force-dependent unfolding of the talin mechanosensitive domain (R3) [talin WT model, see Appendix C, SI]. As for the extracellular part of clutches, it was represented in the model by a linear spring according to the theory of elasticity, see Methods.

Fitting experimental data on the mechanosensitive behaviour of MEF WT cells from ref. [26] to the stationary solution of the model given by Eq. (8)-Eq. (9), using the three model fitting parameters listed in Table 1, revealed that even at the highest substrate stiffness, most talin-based molecular clutches experience relatively low tension in the range of 0.5 − 1.3 pN [see Figure 3(a,b) and S4(b), SI]. This tension is below the force level at which the nonlinear elastic properties of talin become significant [Figure 3(c)], suggesting that unfolding of the mechanosensitive talin domain is potentially not a major factor affecting the mechanical response of adhesion complexes under normal experimental conditions. Consistent with these results, our model predicts that on rigid substrates, only a small percentage (1% − 2%) of talin-based molecular clutches are subject to forces sufficient to substantially stretch talin molecules and trigger the unfolding of their mechanosensitive domain, see Figure S4(a), SI. This finding is in good agreement with a recent experimental study performed on *Xenopus laevis* cells grown on a fibronectin-coated glass substrate, in which only about ∼ 4% of talin molecules underwent significant conformational changes upon molecular clutch stretching [42].

**FIG. 3.**
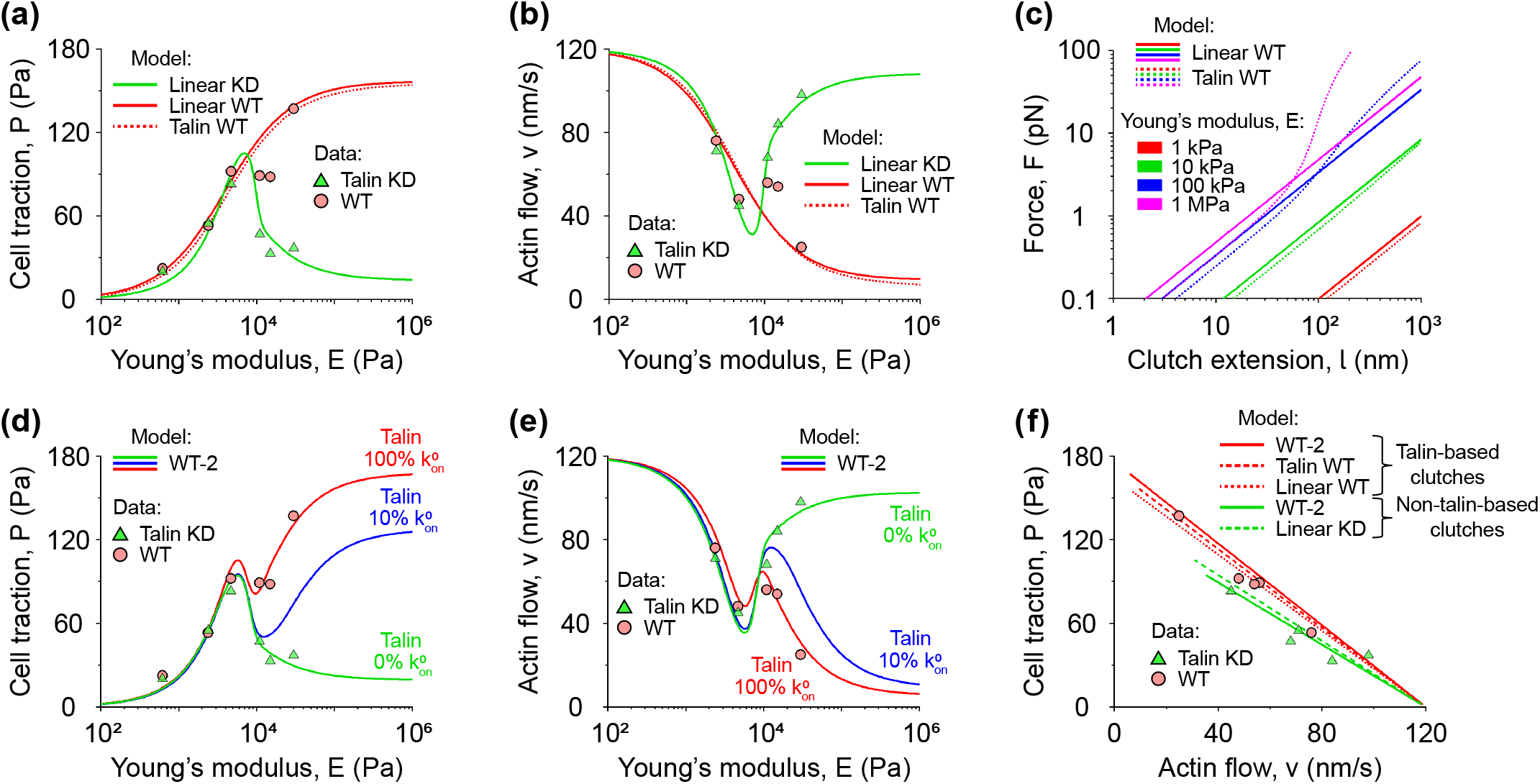
Two types of molecular clutches and the role of local substrate deformations and elastic properties of talin in cell mechanosensing. **(a, b)** Fitting of the cell traction and retrograde actin flow experimental data from ref. [26] collected on MEF WT cells (WT data, red circles) and MEF Talin 1 KO, Talin 2 shRNA cells (talin KD data, green triangles) to the linear clutch model (linear WT and linear KD model, solid curves) and to the nonlinear talin clutch model (talin WT model, dotted curves), respectively. The plot shows that both the linear WT model and the talin WT model provide a fairly accurate fit to the experimental data, suggesting that the approximation of talin molecules by linear springs works quite well. Furthermore, from Table 1 showing the values of the fitting model parameters, it can be seen that there are two types of molecular clutches – soft (talin-based) and rigid, which manifest themselves in MEF WT and MEF Talin 1 KO, Talin 2 shRNA cells, respectively. **(c)** Force-extension curves of linear molecular clutches (linear WT model, solid lines) and nonlinear talin-based molecular clutches (talin WT model, dotted curves) at different values of the substrate Young’s modulus. From the graph it can be seen that the nonlinear properties of talin molecules become apparent only in the case of rigid substrates (*E >* 10 kPa). **(d, e)** Fitting of the cell traction and retrograde actin flow experimental data from ref. [26] collected on MEF WT cells (WT data, red circles) and MEF Talin 1 KO, Talin 2 shRNA cells (talin KD data, green triangles) to the WT-2 model describing competitive formation of the two types of molecular clutches (soft nonlinear talin-based molecular clutches and rigid linear non-talin-based molecular clutches, see Table T2, SI). It can be seen that the WT-2 model more closely matches the experimental data obtained on MEF WT cells compared to the linear WT and talin WT models, see panels (a, b). Moreover, the WT-2 model predicts that even 10% of the maximum density of talin in cell adhesion areas and therefore 10% of the rate of talin-based molecular clutch formation 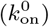 *k*^0^) is sufficient to stabilize cell adhesion to rigid substrates (*E >* 10 kPa), but not to soft substrates (*E <* 10 kPa), see the blue curves. **(f)** Existing experimental data and models tested in this study show a negative linear correlation between cell traction and retrograde actin flow, which arises from the linear force-velocity relationship of myosin II motors, see the Appendix D, SI for details.

These results suggest that the mechanical properties of talin-based molecular clutches in living cells are predominantly determined by their passive elasticity rather than force-dependent conformational changes, while the latter likely contribute to downstream signal mechanotransduction.

Notably, although under normal conditions molecular clutches experience the low average tension mentioned above, variation of the model parameters showed that reduction in the dissociation rate of molecular clutches (*k*_off_) and the density of sites available for the formation of molecular clutches (*σ*_c_) can lead to a significant increase in the average tension of molecular clutches on rigid substrates (to 30 pN), resulting in the force-induced unfolding of the mechanosenstive talin domain in ∼ 70% of the molecules, see Figure S5, SI. This finding suggests that the type of the ECM ligands (collagen / fibronectin / RGD) and integrins interacting with them, as well as the density of the ligands, can strongly influence experimental data and therefore should be precisely controlled in experiments.

### Molecular clutch elasticity is an important factor for adhesion mechanosensing

The talin WT model shows that most molecular clutches experience low tension at which they can be well approximated by linear springs. To test whether this type of passive elasticity is one of the main factors determining mechanosensitive cell adhesion behaviour, we fitted experimental data collected on MEF WT cells and MEF Talin 1 KO, Talin 2 shRNA cells from ref. [26] to a simplified model, in which both the intracellular and extracellular parts of molecular clutches are represented by linear springs. To this aim, we used a few physiologically relevant fitting parameters, including the elasticity of the intracellular part of molecular clutches, listed Table 1 (see the linear WT and linear KD models). From Figures 3(a,b) it can be seen that the linear molecular clutch model was able to fit the experimental data from ref. [26] as accurately as the talin WT model.

The values of the fitting model parameters were found to be in good agreement with previous experimental studies, suggesting that the linear molecular clutch model is able to correctly capture the main physiochemical properties of molecular clutches. For example, the experimental data fitting to the linear molecular clutch model indicates that in the case of MEF WT cells, the elasticity of the intracellular parts of molecular clutches (*k*_c_ = 0.05 pN/nm) is close to the elasticity of talin proteins estimated from previous single-molecule measurements (0.09 pN/nm, see Table T3, SI). Interestingly, the intracellular parts of molecular clutches in MEF Talin 1 KO, Talin 2 shRNA cells are predicted to be more than ten times stiffer (*k*_c_ = 1.6 pN/nm) compared to the case of MEF WT cells, suggesting that cells potentially use at least two different types of molecular clutches to establish adhesion bonds with the substrate – soft (talin-based) and rigid. Moreover, the latter have a more than ten times larger characteristic size of the adhesion site (*r* = 26 nm) compared to talin-based molecular clutches (*r* = 0.6 − 0.8 nm), see Table 1. Because previous single-molecule studies have shown that typical cytoskeletal and FA proteins have elasticities less than 0.3 pN/nm (Table T3, SI), these data suggest that rigid molecular clutches may be formed by clusters of adaptor proteins. It has been shown that more rigid FA proteins, such as kindlin and *α*-actinin, bind to FAs before talin, and *α*-actinin is able to form clusters / condensates [43], which, therefore, may be prime candidates for the main components of molecular clutches in the talin knock out cells.

Importantly, to fit the experimental data, we did not need to assume the force-dependent enhancement of molecular clutches by vinculin, suggesting that vinculin might be more of a downstream factor in cell adhesion mechanics, playing a more prominent role in mechanotransduction rather than force-transmission, in good agreement with previous findings [39–41].

Because both soft talin-based and rigid molecular clutches must be present in MEF WT cells, we next fitted experimental data obtained from such cells to a model describing the competitive formation of the two types of molecular clutches (WT-2 model) – rigid linear molecular clutches corresponding to those found in MEF Talin 1 KO, Talin 2 shRNA cells, and soft nonlinear talin-based molecular clutches. This model was found to more closely match experimental data obtained on MEF WT cells [Figures 3(d,e)], with the values of most model fitting parameters being either identical or very close compared to the case when experimental data were fitted to a single type of molecular clutches, see the linear KD and talin WT models in Table T1 and the WT-2 model in Table T2, SI. Moreover, the value of the spring constant of rigid molecular clutches (*k*_c_ = 0.9 pN/nm) turned out to be very close to the estimates of the rigidity of *α*-actinin clusters (0.8 pN/nm), since each such cluster contains on average four molecules of *α*-actinin [43], each of which has a spring constant of 0.2 pN/nm, see Table T3, SI. This result further supports the model prediction regarding the two types of molecular clutches in MEF WT cells.

Finally, by varying the spring constant of the intracellular part of molecular clutches (*k*_c_) in the linear WT model, it was found that the cell traction curve and the retrograde actin flow curve can undergo significant changes on rigid substrates, see Figure 4. Thus, although most molecular clutches experience low tension, their linear elastic response plays an important role in determining cell adhesion behaviour on rigid substrates.

**FIG. 4.**
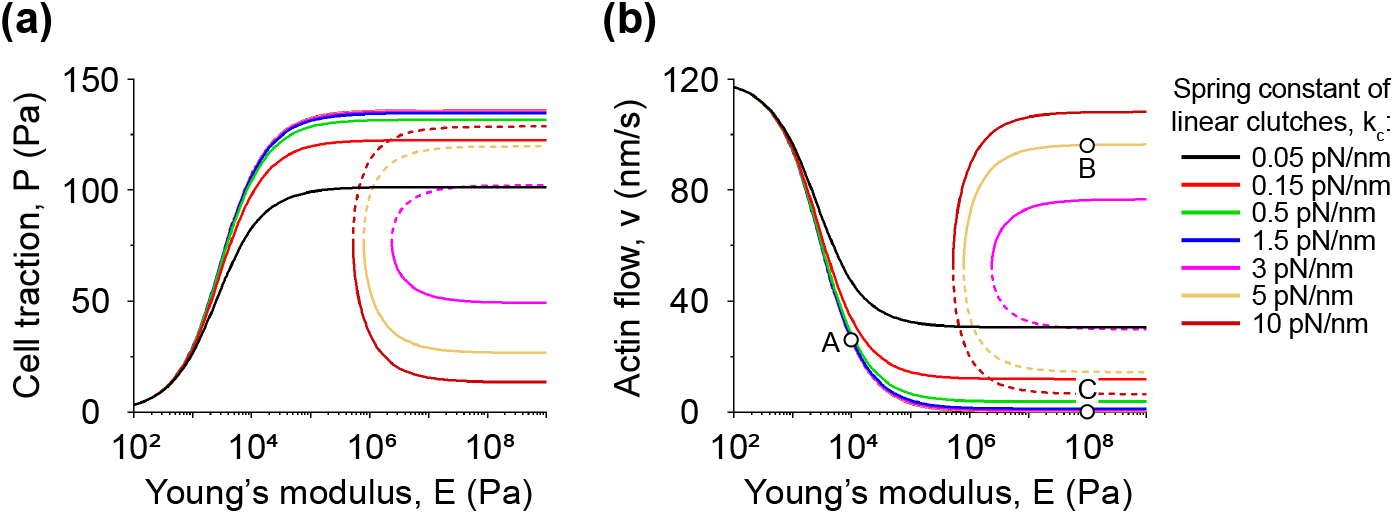
Effect of the elasticity of the intracellular part of molecular clutches on cell adhesion. **(a, b)** Graphs of the cell traction and retrograde actin flow curves calculated using the linear WT model for various values of the spring constant (*k*_c_) describing the elastic properties of the intracellular part of molecular clutches. Solid curves denote stable branches of the graphs, and dashed curves indicate unstable branches, see Figure 2 for details. It can be seen from the plots that the spring constant *k*_c_ has a strong influence on the cell adhesion behaviour in the case of rigid substrates (*E >* 10 kPa), while there is practically no change on soft substrates (*E* ≤ 10 kPa). In addition, the figure shows that in the case of stiff molecular clutches (*k*_c_ ≳ 2.6 pN/nm), the molecular clutch system undergoes bifurcation, leading to bistability of the cell traction and retrograde actin flow curves on rigid substrates. For points A, B and C shown in panel (b), which correspond to the case of *k*_c_ = 5 pN/nm, we performed stochastic simulations and finite-difference calculations of the time evolution of the molecular clutch system by solving the master equation [Eq. (A10), SI], confirming that the molecular clutch system exhibits bistable behaviour on rigid substrates but not on soft ones, see Figure S6(b), SI. To demonstrate the bifurcation, the model parameter values used in the calculations were the same as for the case of low fibronectin density (1*µ*g/ml) shown in Figure 8(a).

### Local substrate deformations are important for the mechanosensitive behaviour of cell adhesion

In addition to molecular clutch elasticity, we also examined the potential role of local substrate deformations at the adhesion sites of molecular clutches in cell mechanosensing. Indeed, experimental data show that mechanical forces can be effectively transmitted from a cell to the substrate by individual molecular clutches [44]. Furthermore, from the theory of elasticity it is known that elastic substrate deformations are inversely proportional to the linear size of the adhesion area to which mechanical stress is applied [34] [Eq. (1)]. Taking into account the experimentally measured characteristic size of FAs (∼ 1.7 *µ*m [26]) and the typical size of the binding sites of adhesion receptors (up to several nanometers), local substrate deformations at the adhesion sites of molecular clutches should be significantly greater than the substrate deformation averaged over the FA scale, and therefore have a much stronger effect on the dissociation kinetics of molecular clutches compared to the average substrate deformation.

Indeed, from Eq. (4), Eq. (5) and Eq. (9) it follows that when the local substrate deformations are completely neglected (i.e., 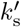 model parameter is set equal to infinity), the sensitivity of the molecular clutch system to the elastic properties of the substrate is significantly reduced – the probability distribution *p*_on_(*l*) of observing a given site occupied by a molecular clutch with extension *l* becomes independent of the Young’s modulus of the substrate, *E*. Thus, in the absence of local substrate deformations at the adhesion sites of molecular clutches, the molecular clutch system loses its mechanosensitive properties after relaxation to the stationary state. This finding is in good agreements with previous experimental studies showing that cells do indeed apply local contractions to the substrate, which serve as a critical step in rigidity sensing [45], highlighting the importance of local substrate deformations by molecular clutches. Moreover, this is consistent with the nature of many biological ECMs, which are heterogeneous in both composition and topology.

Furthermore, comparison of the force-extension curves of linear and nonlinear talin-based molecular clutches from the linear WT model and the talin WT model at different values of the substrate Young’s modulus (*E*) shows that on soft substrates (*E* ≤ 10 kPa) the mechanosensitive behaviour of cell adhesion is determined primarily by the elastic characteristics of the extracellular part of molecular clutches, while on rigid substrates (*E >* 10 kPa) the elasticity of the intracellular part of molecular clutches begins to play a more important role in this process, see Figure 3(c). This result is in good agreement with the graphs shown in Figure 4, which indicate that the spring constant of the intracellular part of molecular clutches (*k*_c_) affects only cell behaviour on rigid substrates, but not as much on soft ones. Thus, elasticity of both intracellular and extracellular parts of molecular clutches is a key factor influencing mechanosensitve cell adhesion behaviour.

### Bifurcation in the molecular clutch system causes bistability of cell adhesion on rigid substrates

The semi-analytical nature of the models developed in our study also allowed for a detailed analysis of the stability of the molecular clutch system and bifurcations experienced by it. It was found that at high values of the spring constant of the intracellular part of molecular clutches (*k*_c_), the cell traction curve and the retrograde actin flow curve often undergo bifurcation, which leads to the appearance of two additional stationary branches – stable and unstable, see, for example, Figure 4.

To map the bistability region in the space of model parameters, we varied the values of *k*_c_ and the myosin II density (*σ*_m_) in the linear KD and linear WT models, see Figure 5. As a result, it was found that in the case of MEF Talin 1 KO, Talin 2 shRNA cells, the bistability region is localized at moderate values of the myosin II density, close to the parameter range of the linear KD model corresponding to MEF Talin 1 KO, Talin 2 shRNA cells [Figure 5(a,e)]. This finding suggests that molecular clutches likely experience bistability in myosin II-depleted regions of such cells that can often be found at the cell periphery [32]. Indeed, fitting of the pre-viously published experimental data collected on MEF Talin 1 KO, Talin 2 shRNA cells treated with myosin II inhibitor, blebbistatin [26], to the linear KD model predicted that these cells begin to develop bistable cell adhesion behaviour when the density of myosin II motors in them is decreases by ≳ 1.6 times [15 *µ*M blebbistatin case, Figure 6(a)], with a further decrease in the myosin II density leading to a more pronounced bistability of cell adhesion [50 *µ*M blebbistatin case, Figure 6(a)]. On the other hand, blebbistatin-treated MEF WT cells do not appear to exhibit bistable cell adhesion behaviour [Figure 6(b)], mainly due to the very high elasticity of talin-based molecular clutches, which results in MEF WT cells being far from the corresponding bistability region in the model parameter space, see Figure 5(b,e). This finding suggests that talin molecules could potentially play an important role in stabilizing nascent adhesions (NAs) during their maturation.

**FIG. 5.**
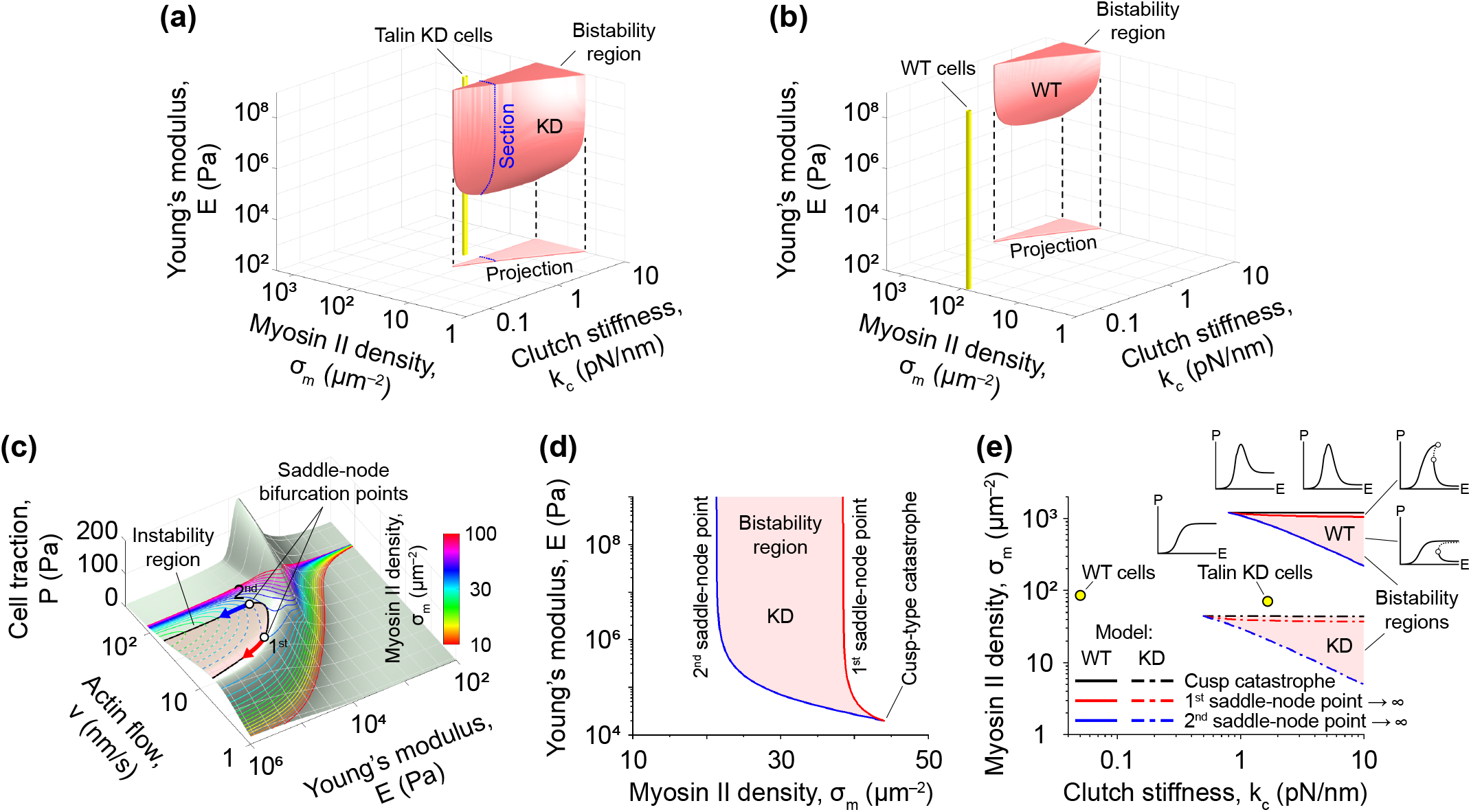
Bistability of cell adhesion. **(a, b)** Stability diagrams of the linear KD and linear WT models, respectively. Red 3D shapes and red triangles indicate the bistability regions and their projection onto the horizontal plane, accordingly. Yellow cylinders represent the range of model parameters corresponding to MEF Talin 1 KO, Talin 2 shRNA cells and MEF WT cells. The blue dotted line in panel (a) indicates a section of the bistability region, which is shown in more detail in panel (d). From panels (a, b) it is clear that while MEF Talin 1 KO, Talin 2 shRNA cells are located close to the bistability region, MEF WT cells are far from it due to the high elasticity of talin-based molecular clutches, suggesting a potential role of talin in stabilizing NAs and FAs. **(c)** Cusp catastrophe experienced by the molecular clutch system. The pale green surface shows cell traction (*P*) as a function of retrograde actin flow (*v*) and Young’s modulus of the substrate (*E*) in the linear KD model. Coloured solid and dashed lines indicate stable and unstable branches of cell traction curves calculated at different myosin II densities (*σ*_m_). The graph shows that a decrease of the myosin II density to ∼ 44 *µ*m^−2^ leads to the appearance of two saddle-node bifurcation points, which is a typical sign of a cusp catastrophe. A further drop in myosin II density leads to the movement of the two saddle-node points along the surface, outlining an instability area that includes only points corresponding to unstable branches of the cell traction curves. **(d)** A section of the bistability region in the linear KD model. Being projected onto the horizontal Young’s modulus axis, the trajectories of the two saddle-node points shown in panel (c) form the boundary of a section of the bistability region as depicted in panel (d), which is the same section as the one indicated in panel (a). Thus, it can be concluded that the bistability region in the model parameter space arises due to a cusp-like catastrophe that the molecular clutch system experiences in the case of a sufficiently high rigidity of the intracellular parts of molecular clutches. **(e)** Projections of the bistability regions shown in panels (a, b) onto the horizontal plane. The graph displays the onset of the cusp catastrophe (black curves) in the linear WT and linear KD models, as well as bifurcations associated with the displacements of the first and second saddle-node points to infinity (red and blue curves, respectively). Yellow dots indicate model parameter values corresponding to MEF Talin 1 KO, Talin 2 shRNA cells and MEF WT cells, accordingly. The insets around the bistability region of the linear WT model show the characteristic shapes of cell traction curves in the corresponding regions of the model parameter space. From the graph presented in panel (e), it can be seen that the bifurcations responsible for the emergence of the bistability region affect not only the shape of the cell traction curves within the region itself, but also outside it.

**FIG. 6.**
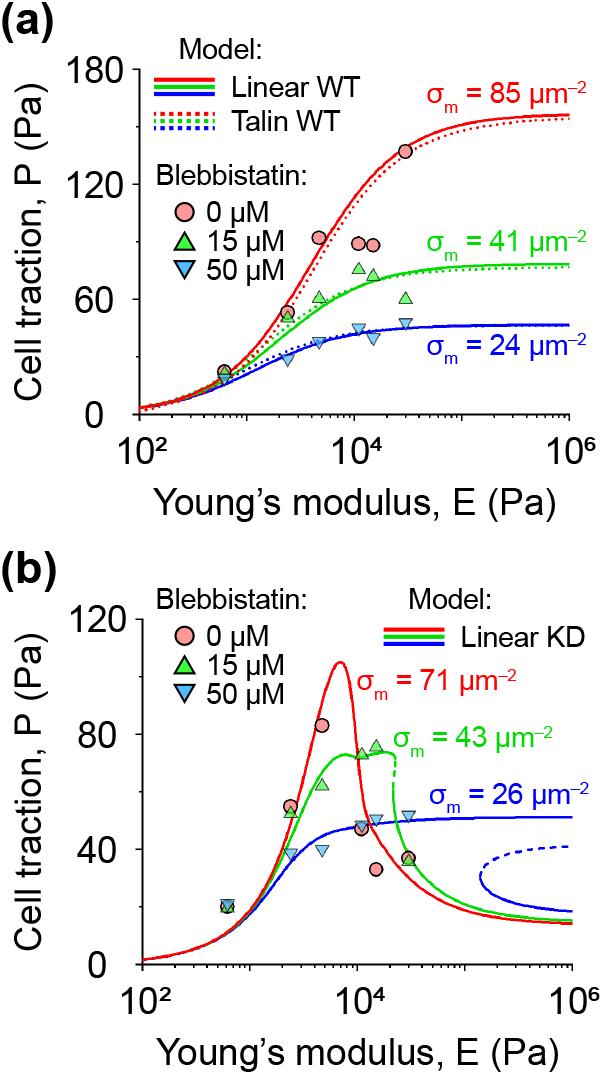
The role of myosin II density in mechanosensitive cell adhesion. **(a, b)** Fitting of experimental data collected on MEF Talin 1 KO, Talin 2 shRNA cells (a) and MEF WT cells (b) treated with different concentrations of the myosin II inhibitor, blebbistatin [26]. The data were fitted to the linear KD model [solid curves, panel (a)], linear WT model [solid curves, panel (b)] and talin WT model [dotted curves, panel (b)], respectively, by varying only the density of myosin II motors. The remaining model parameters were fixed at the values shown in Table T1, SI. The dashed segments of the cell traction curves in panel (a) indicate unstable branches of the corresponding graphs. Panels (a, b) show that while MEF Talin 1 KO, Talin 2 shRNA cells are predicted to have bistable behaviour on rigid substrates at low myosin II densities, MEF WT cells lack this type of behaviour due to the high elasticity of talin-based molecular clutches, see Figure 5 for more details.

To better understand the origin of the bistability of cell adhesion and the potential role of myosin II density (*σ*_m_), we varied the latter in the linear KD model, plotting cell traction (*P*) as a function of retrograde actin flow (*v*) and Young’s modulus of the substrate (*E*), see Figure 5(c). It was found that a decrease in the myosin II density to 44 *µ*m^−2^ leads to the appearance of an unstable branch with two saddle-node bifurcation points at its ends, where the unstable branch merges with the stable ones, – a typical sign of a cusp catastrophe [Figure 5(c)]. A further decrease in the myosin II density in the linear KD model led to the displacement of the two saddle-node points along the surface describing cell traction (*P*), outlining a region of instability that includes only points corresponding to the unstable branches of the cell traction curves shown in Figure 5(c). Being projected onto the horizontal Young’s modulus axis, the trajectories of the two saddle-node points form the boundary of a section of the bistability domain [Figures 5(a,d)]. Thus, it can be concluded that the bistability region arises in the model parameter space due to a cusp-like catastrophe that the molecular clutch system experiences in the case of a sufficiently high rigidity of the intracellular part of molecular clutches.

Finally, it should be noted that the catch-bond behaviour of adhesion receptors (integrins) turned out to be dispensable for the occurrence of all the above bifurcations, since very similar behaviour of the molecular clutch system was also observed in the case of slip-bonds [Eq. (11) and Eq. (12)], suggesting that the bistable behaviour is an intrinsic property of the system.

### Cell adhesion complexes function close to a stationary / quasi-stationary state

The existence of stable stationary states at each value of the substrate Young’s modulus suggests that the molecular clutch system should converge over time to one of these states. To find out how quickly the molecular clutch system reaches these stationary states, starting from an initial configuration with zero adhesion bonds between a cell and the substrate, we used the finite-difference method to numerically solve the master equation describing the time evolution of the molecular clutch system [Eq. (A10), SI]. As a result, both the linear WT and linear KD models were found to predict rapid convergence of key experimental observables, such as cell traction and retrograde actin flow, to their stationary values within 10 − 20 s, see Figures 7(a,b) and S7(a,b), SI. The probability distribution of molecular clutch extension also demonstrated rapid convergence to the stationary distribution given by Eq. (9) in as little as 20 s [Figures 7(c) and S7(c), SI].

**FIG. 7.**
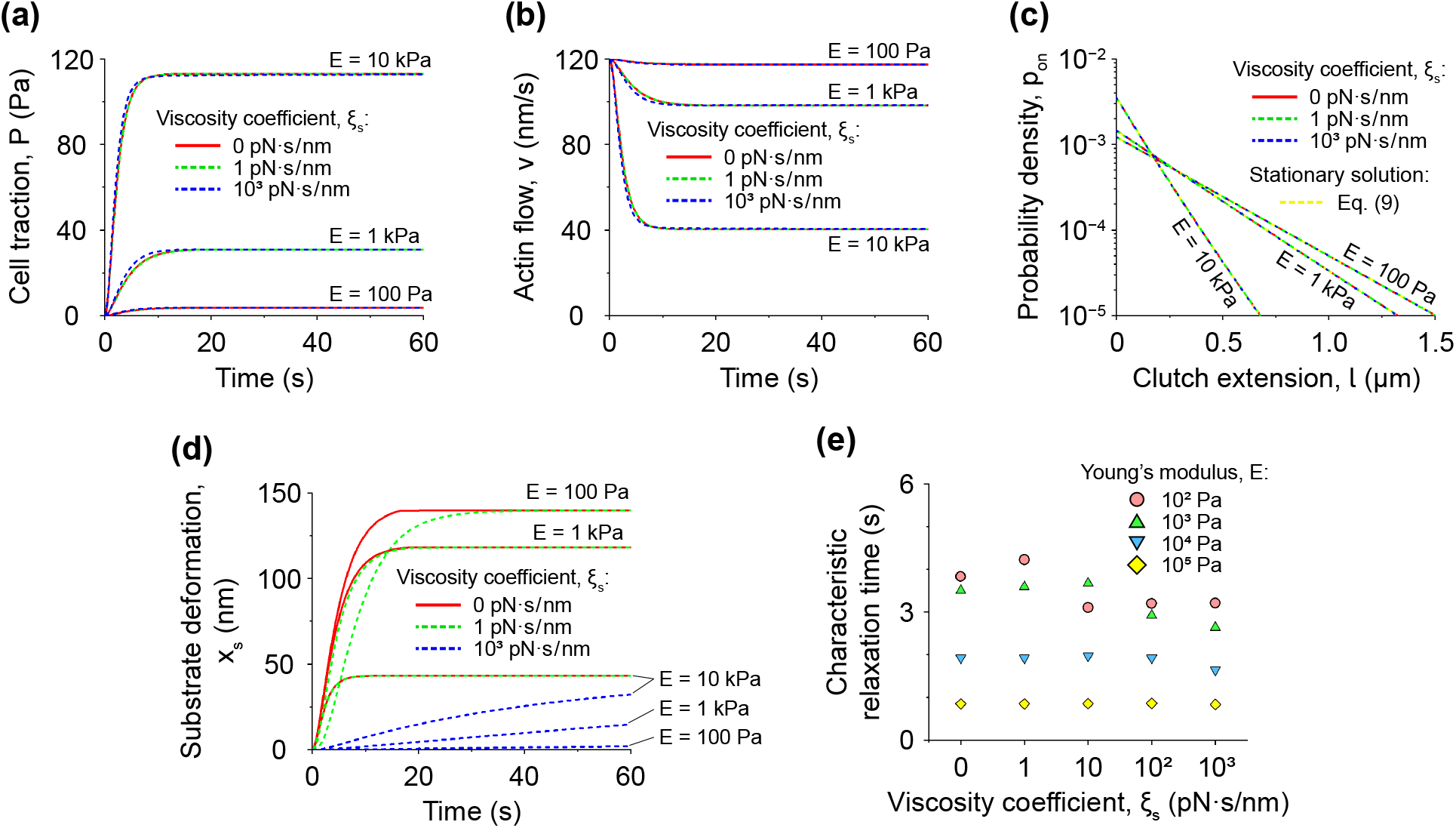
Time evolution of cell adhesion in MEF WT cells. **(a, b)** Time evolution of cell traction (*P*), retrograde actin flow (*v*) and substrate deformation (*x*_s_), starting from an initial configuration with zero molecular clutches between cell and substrate. The curves were obtained by numerically solving the master equation describing the time evolution of the system [Eq. (A10), SI] using the finite-difference method. Calculations were carried out for three different values of the substrate Young’s modulus (*E*): 100 Pa, 1 kPa and 10 kPa. The substrate viscosity coefficient (*ξ*_s_) in the same calculations was varied from 0 pN · s/nm to 1000 pN · s/nm. The values of the remaining model parameters were the same as those in the linear WT model, see Table T1, SI. It can be seen that cell traction and retrograde actin flow quickly reach stationary values within ∼ 10 − 20 s, regardless of the substrate viscosity. **(c)** Probability distribution (*p*_on_) of molecular clutch extension (*l*), calculated after 20 s time evolution of the molecular clutch system for three different values of Young’s modulus (*E*) and viscosity coefficient of the substrate (*ξ*_s_). The graph shows that the molecular clutch system reaches the stationary / quasi-stationary probability distribution described by Eq. (9) (yellow dashed lines) within 20 s, regardless of the substrate viscosity. **(d)** Time evolution of substrate deformation. As can be seen from the graph, the characteristic relaxation time of the substrate is highly dependent on its viscosity, ranging from a few seconds to many minutes. Yet, from panels (a-c) it is clear that slow substrate relaxation has little effect on the rapid convergence of the probability distribution of molecular clutch extension, as well as cell traction and actin retrograde flow to their stationary / quasi-stationary values / states. **(e)** Dependence of the characteristic relaxation time of retrograde actin flow on Young’s modulus and viscosity coefficient of the substrate. The characteristic relaxation time was calculated by fitting the curves shown in panel (b) to an exponential decay function. As can be seen from the graph, the characteristic relaxation time decreases with increasing Young’s modulus of the substrate. However, it is practically independent of the viscous properties of the substrate.

These results are in good agreement with previous experimental observations indicating that retrograde flow of actin filaments in filopodia or near the edges of living cells occurs at an almost constant rate (*v*(*t*) ≈ const) [20, 26, 31, 32], suggesting that cell adhesion complexes should function close to a steady-state. On the other hand, these results are in sharp contrast to the predictions of the conventional molecular clutch theory, which suggests that retrograde actin flow should experience strong fluctuations under similar conditions [27], also see Figure S6(a), SI. Thus, our model better reflects existing experimental data. Moreover, the average density of engaged molecular clutches predicted by the linear WT, talin WT and WT-2 models in the case of MEF WT cells was found to be relatively high [50 − 150 *µ*m^−2^, Figure S4(a,c), SI] compared to previous models (≲ 10 *µ*m^−2^ [20, 24, 27]), which is also in better agreement with experimental estimations of the talin density in FAs (∼ 80 − 500 *µ*m^−2^ [43]). Furthermore, the loading rates of molecular clutches in the stationary state predicted by the linear WT, talin WT and WT-2 models in the case of rigid substrates (0.2 − 5.5 pN/s, *E* = 1 MPa, Figure S8, SI) are also in better agreement with recent experimental studies (0.5 − 4 pN/s [33]), in sharp contrast to the predictions made by previous molecular clutch models (*>* 100 pN/s [20, 24–27]).

Interestingly, by fitting the calculated curves of retrograde actin flow to an exponential decay function, it was found that the average value of the characteristic relaxation time of the molecular clutch system (*τ*_relax_ = 0 − 4 s) is practically independent of the viscous properties of the substrate, although still dependent on its Young’s modulus, see Figures 7(e) and S7(e), SI. In addition, it turned out that in the case of viscous substrates (*ξ*_s_ ≥ 100 pN · s/nm), the cell traction and retrograde actin flow curves reach their quasi-stationary values within the same 10 − 20 s, despite the fact that the substrate deformation continues to change slowly with time [Figures 7(d) and S7(d)]. This result suggests that in the case of viscoelastic materials described by the Kelvin-Voigt model, which was used in our model to represent large-scale substrate deformations [Figure 1(b)], the viscous properties of the substrate have little effect on the rapid establishment of a stationary / quasi-stationary state by cell adhesion complexes.

Furthermore, it can be seen from Table T4, SI that the predicted relaxation time of the molecular clutch system is close to the characteristic dissociation time of talin-based molecular clutches measured in experiments, suggesting that our model provides physically reasonable estimates.

Notably, the relaxation time predicted by our model turns out to be 2 − 4 orders of magnitude less than the average lifetime of FAs [Table T4, SI], indicating that they spend more than 98% of their time in a stationary / quasi-stationary state. This finding is in good agreement with the above conclusions regarding the steady-state behaviour of cell adhesion complexes based on experimental observations of the nearly constant rate of retrograde flow of actin filaments associated with these complexes.

In addition, from Table T4, SI it is clear that the experimentally measured characteristic binding and dissociation times of vinculin and paxillin to / from FAs are an order of magnitude greater than the characteristic relaxation time of the molecular clutch system. This result suggests that talin may serve as a rapid response protein that helps establish the initial landscape of molecular clutches in a substrate-mechanosensitive manner, which can subsequently be reinforced and shaped by vinculin / paxillin into matured FAs, in good agreement with previous experimental data [10, 26, 39–41].

### Ligand concentration regulates cell adhesion via a myosin II feedback loop in fibroblast cells

Finally, to investigate the effect of the density of molecular clutches on cell adhesion, we fitted previously published experimental data on MEF WT cells and MEF Talin 1 KO, Talin 2 shRNA cells plated on elastic substrates that were coated with different fibronectin concentrations [26]. Since the density of fibronectin network affects the rate of molecular clutch formation, as well as the distribution of mechanical load near adhesion sites of molecular clutches, the data fitting was performed by varying the clutch formation rate and the characteristic radius of adhesion sites of molecular clutches. In addition, the density of myosin II motors also varied. Using this approach, the experimental data could be fitted fairly well to the the linear KD, linear WT and talin WT models, see Figure 8(a,b).

**FIG. 8.**
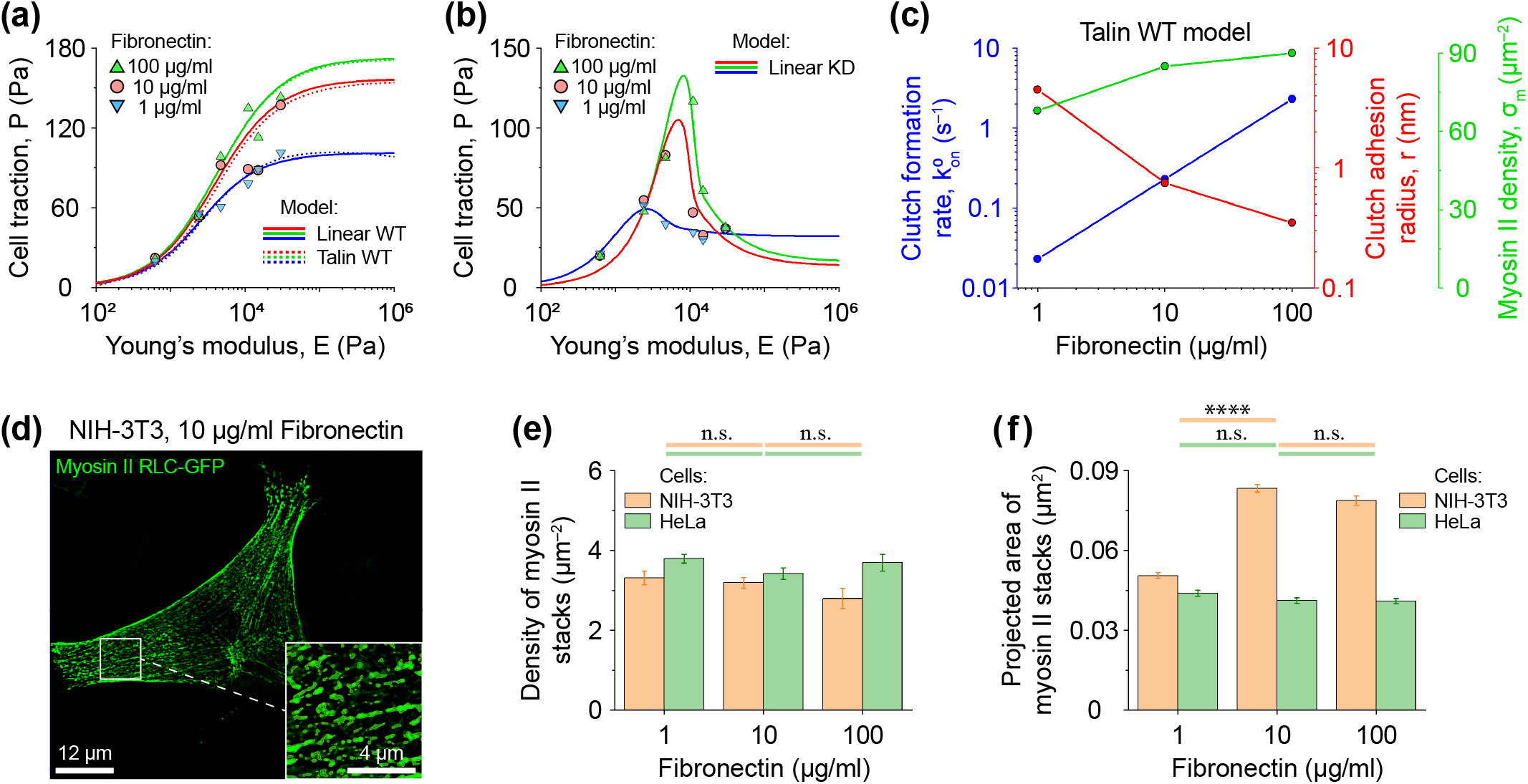
Effect of fibronectin concentration on cell adhesion. **(a, b)** Fitting of experimental data collected on MEF WT cells (a) and MEF Talin 1 KO, Talin 2 shRNA cells (b) at different concentrations of fibronectin solution used to coat the substrate surface [26]. The data were fitted to the talin WT model (dotted curves), and the linear WT and linear KD models (solid curves) by varying the rate of molecular clutch formation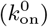, the radius of adhesion sites of molecular clutches (*r*), and the density of myosin II motors (*σ*_m_). The rest of the model parameters were fixed at the values given in Table T1, SI. **(c)** Dependence of the values of the fitted parameters in the talin WT model on the concentration of fibronectin. From the graph it can be seen that the model predicts that higher fibronectin concentration leads not only to faster molecular clutch formation, but also to a decrease in the effective radius of adhesion sites of molecular clutches. Furthermore, a higher concentration of fibronectin results in an increase in the density of myosin II motors, which exert mechanical load on the actin cytoskeleton. **(d)** SIM image of an NIH-3T3 fibroblast cell expressing myosin RLC-GFP construct. The inset shows a magnified area indicated by the white square in the figure. **(e, f)** Measured density (e) and projected area (f) of RLC-GFP-labelled myosin II stacks in NIH-3T3 and Hela cells as a function of the fibronectin concentration used to coat glass slides. Each column shows the mean calculated from *N* = 834 − 3836 myosin II stacks from 11 − 16 cells, errorbars represent SEM. The bars above the graphs indicate the pairwise statistical difference between the corresponding columns (n.s. – non-significant, **** – *p <* 0.0001). Panels (e, f) demonstrate that although the density of myosin II stacks was independent of fibronectin concentration in the tested cell lines, their average projected area varied with fibronectin concentration depending on cell type. In particular, NIH-3T3 cells show qualitatively the same trend as predicted by the talin WT model for MEF WT cells.

Interestingly, the data fitting showed a strong correlation between myosin II motor density (*σ*_m_) and fibronectin concentration as the fitting did not converge when myosin II density was fixed. According to the model, the density of myosin II motors pulling on actin filaments decreases by ∼ 25% on substrates coated with 1 *µ*g/ml fibronectin solution compared to the case of 100 *µ*g/ml fibronectin solution, see Figure 8(c). To test this model prediction, we transfected NIH-3T3 cells with myosin regulatory light chain (RLC)-GFP [46] and measured the density and the average size of myosin II stacks labelled with this construct as a function of the fibronectin concentration, see Figures 8(d-f). As a control, we also performed similar measurements in HeLa cells.

It was found that while in HeLa cells myosin II motors formed more or less uniform assemblies resembling beads on a string [Figure S9, SI], in NIH-3T3 cells they formed more irregular stacks, see Figure 8(d). More importantly, although the fibronectin concentration was found to have little effect on the density of myosin II stacks in both NIH-3T3 and HeLa cells [Figure 8(e)], it had a strong effect on their average size in NIH-3T3 cells (∼ 35% drop, 1 *µ*g/ml fibronectin solution compared to 100 *µ*g/ml solution), but not in HeLa cells [Figure 8(f)], suggesting that this effect may be cell-type dependent. Thus, our model can quite accurately capture the mechanical states of fibroblast cells and make physiologically relevant predictions.

Previous studies have shown that fibronectin density can influence the activation of Rho kinase (ROCK) in endothelial cells, which controls the assembly of myosin II stacks [47]. Thus, MEF cells could potentially use a similar mechanism. However, the precise molecular pathways responsible for fibronectin-dependent activation of ROCK remain unknown. Since our observations indicate that such a feedback loop is cell type-dependent, it may be possible in the future to gain detailed insight into the origin of this phenomenon by comparing normal and transformed cells.

## DISCUSSION

In this work, we developed a semi-analytical model of cell adhesion based on the molecular clutch theory to examine the potential roles of adaptor protein elasticity, local substrate deformations, myosin II and ligand densities on mechanosensitive cell adhesion behaviour.

The main difference between our model and previous studies is that we took a more explicit approach to modeling the elastic properties of adaptor proteins and substrate deformations. Then, by deriving a semi-analytical solution to the differential master equation of the model, we were able to evaluate experimental observables characterizing the mechanosensitive behaviour of cell adhesion complexes, such as cell traction force and the rate of retrograde actin flow, using experimentally measured parameters such as the stiffness of adaptor proteins and the dissociation rate of molecular clutches. Based on experimental data fitting, performed by varying a few key model parameters (Table 1), our model was able to accurately reproduce all previous experimental observations and make predictions regarding the correlation between ECM ligand and myosin II densities that were successfully tested in experiments. Importantly, the physical picture offered by our model differs from the conventional molecular clutch theory in several essential aspects.

First, our model predicts that the molecular clutch system quickly reaches a stationary state at any substrate rigidity in less than 10 − 15 s. This prediction better reflects the experimentally observed steady retrograde actin flow in living cells under both low and high cell traction conditions [20, 26, 31, 32], which is in sharp contrast to the strong fluctuations predicted by previous models [27]. This means that the mechanodependent dynamics of cell adhesion complexes is a fast and robust process that allows easy differentiation of soft and rigid substrates without having to go through multiple stochastic load-and-fail cycles, which can lead to an increase in the number of missense events.

The predicted characteristic relaxation time of the molecular clutch system to a stationary state (0.1 − 4 s) turned out to be an order of magnitude less than the average turnover time of talin and integrin in FAs (15 − 50 s) and the characteristic binding time of vinculin and paxillin to FAs (30 − 60 s) measured in experiments (Table T4, SI). This result suggests that the dynamic evolution of cell adhesion is a quasi-stationary process mainly driven by talin-based molecular clutches, which are shaped by vinculin and paxillin into higher-order structures such as FAs, in good agreement with previous experimental studies [10].

Our model further highlights the role of the elastic properties of adaptor proteins that constitute molecular clutches, such as *α*-actinin and talin. Namely, the model predicts that the high elasticity of long talin molecules allows them to robustly form molecular clutches on both soft and rigid substrates, in contrast to the previously described load-and-fail mechanism of molecular clutch functioning on rigid substrates [20, 24, 27]. Upon increase of the stiffness of adaptor proteins, – a situation reminiscent of talin knockout / knockdown condition, our model predicts a strong decrease in the density of molecular clutches and the cell traction force. This results in altered cell behaviour, which is in good agreement with experimental studies [26], highlighting the importance of molecular clutch elasticity in governing cell adhesion behaviour.

It should be noted that in our study we considered only the unfolding of the mechanosensitive R3 domain of talin. Yet, it has previously been suggested that mechanical unfolding of other domains may help stabilize the tension experienced by talin, which in turn influences the stability of the corresponding molecular clutches, – an effect that has also been proposed to contribute to the function of many adaptor proteins found in various cells [48, 49]. Moreover, FAs contain many different types of adaptor proteins that have different elastic properties than talin. How the unfolding of multiple talin domains, as well as how mixtures of different molecular clutch species regulate molecular clutch behaviour, is an important question. In the future, combining our model with atomic-level simulations of the stability of globular domains of adaptor proteins [50] may lead to a better understanding of how the protein composition of molecular clutches influences mechanosensitive cell adhesion behaviour in a context-dependent manner.

Notably, several physiological-relevant parameters coming out of our model predictions also appeared to be more in line with experimentally measured values compared to previous theoretical studies. For example, the force loading rates of clutch molecules predicted by our models in the case of rigid substrates (0.2 − 5.5 pN/s, *E* = 1 MPa) are in better agreement with recent direct experimental measurements using single-molecule DNA tension sensors (0.5 − 4 pN/s, [33]) compared to the predictions made by previous molecular clutch models (*>* 100 pN/s [20, 24–27]). Furthermore, the models developed in our work also suggest that talin-based molecular clutches transmit most of the cell traction force on rigid substrates, – in good agreement with recent experimental measurements based on FRET talin tension sensors showing that talin is the main force-bearing molecule in cell adhesion complexes [39, 40], which is in sharp contrast to previous studies predicting that talin-based molecular clutches transmit only ∼ 7% of the cell traction force [26]. These results demonstrate that the theoretical framework developed in our study is capable of accurately relating the force-response of molecular clutches at the single-molecule level to the large-scale ensemble behaviour of cell adhesions.

Our model also predicts that at high adaptor protein and substrate stiffness (*k*_c_ ≳ 0.5 − 0.8 pN/nm, *E* ≳ 100 kPa), the molecular clutch system exhibits bistable behaviour that, from a physical perspective, resembles the cytoadhesion properties of malaria-infected red blood cells [51, 52]. This bistability includes two stable states corresponding to weak and strong cell adhesion, with the former characterized by lower density of molecular clutches, smaller cell traction and faster retrograde actin flow compared to the latter. It was found that the global bifurcations responsible for the emergence of such bistable behaviour determine not only the shape of the cell traction and retrograde actin flow curves within the bistability region, but also far beyond it, resulting in a very distinct mechanosensitive response of WT MEF and MEF Talin 1 KO, Talin 2 shRNA cells to the substrate stiffness [26]. In the future, detailed observations of retrograde actin flow / traction force in talin-depleted cells may shade further light on the fraction and transition dynamics between the two stable states of cell adhesion. The bistability of the molecular clutch system may potentially occur at a single cell level or the level of individual FAs, depending on the heterogeneity of the density and topography of the ECM ligand network with which cell interacts. Indeed, both can influence the local density of sites available for molecular clutch formation and hence the rate of molecular clutch assembly, which was found in our study to have a significant effect on the location of the bistability region of cell adhesion in the model parameter space and, consequently, on the shape of the cell traction and retrograde actin flow curves. This, in turn, can strongly influence the tension experienced by molecular clutches and actin filaments in different cell parts, altering the mechanodependent cooperative as-sembly of myosin IIA and myosin IIB stacks, which has been previously found in experiments [53–55]. The assembly of such stacks is regulated by ROCK, MLCK [55] and PKCz [54] kinases, which are responsible for phosphorylation of the regulatory light chain and heavy chain of myosin IIB, respectively. And while the accumulation kinetics for myosin IIA,C were found to be nearly identical across cell types, myosin IIB exhibited highly cell-type- and cell cycle stage-specific behaviour due to PKCz activity [54]. Thus, taken together, the above results suggest that the heterogeneity and topography of the ECM ligand network should have a significant impact on cell adhesion behaviour, which may be highly cell-type dependent, in good agreement with previous experimental studies [2].

Our model calculations indicate that an increase in myosin II density, which can be caused by the mechanodependent cooperative assembly of myosin IIA and myosin IIB stacks around NAs, can trigger the transition of such NAs from a weak to a strong adhesion state. This, in turn, may lead to talin recruitment and further maturation of NAs, as talin binding to actin has previously been shown to be enhanced by applied mechanical load [36]. Thus, the fibronectin-myosin II feedback loop observed in our study may play an essential role in the maturation of cell adhesion complexes.

Indeed, it was previously shown that inhibition of myosin II in HeLa cells by treatment with blebbistatin or Y-compound, as well as knockdown of the myosin IIA heavy chain, leads to a significant weakening of the filopodia adhesion to fibronectin-coated substrates [32]. Very similar behaviour was also observed in filopodia that lack myosin II at their base region [32]. Since filopodia, which guide cell migration, are often used by cells to probe the local microenvironment and form NAs, these results suggest that the bistable behaviour of molecular clutches could potentially be employed by living cells to move through the ECM or on flat substrates.

Furthermore, our results indicate that the difference between the elasticity of fibronectin and RGD ligands, which are commonly used in mechanobiological studies, and the way these ligands are attached to the underlying substrate (non-specific adsorption or covalent binding) can have a strong effect on the ability of the ligand network to transfer force from cell adhesion complexes to the substrate. This in turn will result in very distinct cell adhesion behaviour.

For example, our model predicts that to probe the elastic properties of the substrate, cells locally apply significant mechanical load to the ECM, pinching it at the adhesion sites of molecular clutches. This result is in good agreement with previous experimental studies performed using elastic micropillar arrays, which showed that cells indeed apply local contractions to the substrate to sense its mechanical properties [45]. On the other hand, it suggests that cell-generated forces can significantly perturb the local organization of the ECM network. In the case of ECM ligands adsorbed to the substrate via non-specific interactions, these local perturbations may eventually accumulate into large-scale changes in the ECM network at the cell site. Indeed, it was experimentally discovered that in this case cells apply a force strong enough to remove surface-adsorbed fibronectin, reorganizing it into fibrils [56]. This, in turn, leads to a transition of cells from FAs to elongated fibrillar adhesions, radically changing the cell adhesion pattern [56].

These differences may be important for understanding the microenvironment sensing behaviour of cells beyond substrate stiffness. Namely, although various materials such as collagen network, or PDMS and glass coated with ECM ligands may have similar bulk elasticity, the behaviour of cells on such substrates may differ significantly due to their different elastic response at the molecular level. In the future, increasingly realistic models of the substrate-molecular clutch coupling may help resolve numerous experimental discrepancies due to such differences and provide valuable information for understanding cell-ECM interactions, which is important for tissue engineering.

It should be noted that one critical missing link for the development of molecular clutch models is how the dynamic behaviour of cell adhesion complexes could potentially be involved in the regulation of downstream cell fate decisions. Many mechanosensitive signaling events involve force-dependent conformational changes of molecular clutch components such as talin, p130cas and FAK, which can alter molecule interaction patterns, enzyme kinetics, and calcium influx. With more accurate and realistic modeling of the conformational states of molecular clutch proteins, which is possible within our model, one can begin to quantitatively investigate the complex interactions between mechanical and biochemical processes at the molecular level that have important consequences for many physiological processes, from morphogenesis to cancer progression.

For example, the quasi-static nature of molecular clutch kinetics and the appearance of bistable cell adhesion behaviour in our model suggest that, for a given substrate rigidity, multiple stable states of molecular clutches can coexist. This means that our model can, in principle, be used as a theoretical basis to describe the maturation process of FAs, in which small NAs gradually transform into mature FAs with a different molecular composition, molecular clutch density and traction force, despite the same extracellular environment. In this context, the maturation of FAs can be viewed as a series of quasi-stationary states, each of which represents a snap-shot of maturing cell adhesion. Then changes in physiological parameters describing mechanical or biochemical processes, such as molecular clutch density / elasticity and myosin II contractility, will collectively model the evolution of quasi-stationary behaviour of FAs towards more mature states. In addition, the bistability of cell adhesion found in our work could also explain why only a small fraction of NAs mature on rigid substrates while the rest disassemble, warranting future studies.

Finally, in addition to substrate stiffness, rheological properties of the ECM, such as viscoelasticity, have also been shown to be an important regulator of cellular functions and are related to pathological processes of tumor growth and metastasis. There have been several interesting attempts to model the mechanosensitive behaviour of cells on viscoelastic substrates based on the conventional molecular clutch theory, which have successfully reproduced cellular behaviours such as periodic oscillations in cell spreading and cell migration [23, 27–30]. In the future, it will be interesting to combine our semi-analytic model with these previous studies to explore how physiological factors, such as the stiffness of molecular clutch molecules, regulate these important cell behaviours.

## AUTHOR CONTRIBUTIONS

P.L. performed experiments. P.L., Q.W. and A.K.E. analysed the data. M.Y. and A.K.E. designed the research and wrote the paper. A.K.E. supervised the study,derived formulas and carried out computations.

## ACKNOWLEDGEMENT

We greatly appreciate the encouraging and insightful discussions with Dr. Jie Yan (MBI, Singapore) and Dr. Alexander D. Bershadsky (MBI, Singapore) about the results of our study. We are grateful to Dr. Alexander D. Bershadsky for providing genetic constructs used in the study. We would like to thank Dr. Qin Peng (Shenzhen Bay Laboratory, China) for NIH-3T3 and HeLa cell lines. Also, we would like to thank Shenzhen Bay Laboratory Supercomputing Center for assistance in carrying out the calculations and Bioimaging Core of Shenzhen Bay Laboratory for providing cell imaging support. We are also grateful to Bioimaging Core engineer Yu Mei for assistance with SIM microscopy (Elyra7). This work was supported by the start-up funds from Shenzhen Bay Laboratory (A.K.E.).

