## Supplementary Information for "Multistable mechanosensitive behavior of cell adhesion driven by actomyosin contractility and elastic properties of force-transmitting linkages"

(Dated: June 14, 2024)

### SUPPLEMENTARY INFORMATION

#### Appendix A: Derivation of the master equation

This section demonstrates the derivation of a closed system of equations describing the dynamics of molecular clutches and the time evolution of $p_{\mathrm{on}}(l,t)$ probability distribution for the linear molecular clutch model discussed in the Methods section of the main text.

To this aim, it is first important to know how quickly the extension of a molecular clutch changes over time due to the retrograde actin flow ($v$) and substrate deformation ($x_{\mathrm{s}}$). By definition, the time-dependent extension of a molecular clutch along the x-axis, $l(t)$, is equal to:

$$l(t) = x_2(t) - x_1(t) \tag{A1}$$

Where $x_1(t)$ and $x_2(t)$ are the x-coordinates of the molecular clutch ends [Figure S2(a)].

If $x_0$ is the coordinate of the initial attachment point of the molecular clutch to the substrate at time $t_0$ [Figure S2(a)], then the x-coordinates of the molecular clutch ends can be found as:

$$\begin{cases} x_1(t) = x_0 + x_{\mathrm{s}}(t) \\ x_2(t) = x_0 + x_{\mathrm{s}}(t_0) + \displaystyle\int_{t_0}^{t} v(t')\,\mathrm{d}t' \end{cases} \tag{A2}$$

Here $x_{\mathrm{s}}(t)$ and $v(t)$ are the average mechanical deformation of the substrate and retrograde actin flow as a function of time.

Differentiating Eq. (A1)-Eq. (A2) with respect to time, it is easy to see that the rate of elongation of the molecular clutch along the x-axis, $v_l(t)$, equals to:

$$v_l(t) = \frac{\partial l}{\partial t} = v(t) - \frac{\partial x_{\mathrm{s}}(t)}{\partial t} \tag{A3}$$

Given Eq. (A3), it is straightforward to derive the master equation describing the time evolution of $p_{\mathrm{on}}(l,t)$ probability distribution of molecular clutch extension. To do this, one needs to divide the axis showing all possible extensions of molecular clutches into infinitesimal intervals of $\mathrm{d}l$ length, as schematically depicted in Figure S2(b), such that the probability distribution $p_{\mathrm{on}}(l,t)$ can be considered uniform on each of these intervals. Then the net change $\mathrm{d}p_{\mathrm{on}}(l,t)$ in $p_{\mathrm{on}}(l,t)$ probability distribution at each of these intervals over a small time step $\mathrm{d}t$ will be the result of the formation / disassembly kinetics of molecular clutches, as well as their stretching due to the retrograde flow of actin filaments and substrate deformation:

$$\mathrm{d}p_{\mathrm{on}}(l,t) = \mathrm{d}p_{\mathrm{on}}(l,t)|_{\mathrm{stretch}} + \mathrm{d}p_{\mathrm{on}}(l,t)|_{\mathrm{kinetics}} \tag{A4}$$

The first term in Eq. (A4) can be obtain simply by considering the flow of probability, which is described in Figure S2(b):

$$\mathrm{d}p_{\mathrm{on}}|_{\mathrm{stretch}}(l,t) = \frac{p_{\mathrm{on}}(l-\mathrm{d}l,t) - p_{\mathrm{on}}(l,t)}{\mathrm{d}l}\, v_l(t)\,\mathrm{d}t = -v_l(t)\,\frac{\partial p_{\mathrm{on}}(l,t)}{\partial l}\,\mathrm{d}t \tag{A5}$$

As for $\mathrm{d}p_{\mathrm{on}}|_{\mathrm{kinetics}}(l,t)$ term, then taking into account the first-order kinetic rates of formation and disassembly of molecular clutches, $k_{\mathrm{on}}(l)$ and $k_{\mathrm{off}}(l)$, which, generally speaking, depend on the extension of molecular clutches, we have:

$$\mathrm{d}p_{\mathrm{on}}|_{\mathrm{kinetics}} = -k_{\mathrm{off}}(l)\,p_{\mathrm{on}}(l,t)\,\mathrm{d}t + k_{\mathrm{on}}(l)\,p_{\mathrm{off}}(t)\,\mathrm{d}t \tag{A6}$$

Where $p_{\rm off}(t)$ is the time-dependent probability of the off-state of molecular clutch sites.

By combining together Eq. (A3)-Eq. (A6), we finally obtain the following formula for the time-dependent evolution of $p_{\rm on}(l,t)$ probability distribution, reminiscent of those described in previous studies [1–3]:

$$\frac{\partial p_{\rm on}(l,t)}{\partial t} = -k_{\rm off}(l)\, p_{\rm on}(l,t) + k_{\rm on}(l)\, p_{\rm off}(t) - \left[v(t) - \frac{\partial x_{\rm s}(t)}{\partial t}\right] \frac{\partial p_{\rm on}(l,t)}{\partial l} \tag{A7}$$

To derive a similar evolution equation for $p_{\rm off}(t)$ probability, all we need to do is to apply $\int_{-\infty}^{+\infty} {\rm d}l$ integration to Eq. (A7) and then use Eq. (2) from the main text. As a result, it can be shown that:

$$\frac{\partial p_{\rm off}(t)}{\partial t} = \int_{-\infty}^{+\infty} \left[k_{\rm off}(l)\, p_{\rm on}(l,t) - k_{\rm on}(l)\, p_{\rm off}(t)\right] {\rm d}l \tag{A8}$$

Where it has been taken into account that $p_{\rm on}(-\infty,t) = p_{\rm on}(+\infty,t) = 0$, i.e., there are no infinitely stretched molecular clutches, since they dissociate long before their extension reaches very large values.

Looking at Eq. (A7), it is easy to see that to get a closed system of equations describing the dynamics of molecular clutches, we still need mathematical formulas for the velocity of the retrograde actin flow, $v(t)$, and the derivative of the average substrate deformation with respect to time, $\partial x_{\rm s}/\partial t$. The former can be obtained by combining Eq. (6) and Eq. (7) from the main text. As for $\partial x_{\rm s}/\partial t$ derivative, from the Kelvin–Voigt model of viscoelastic materials, in which viscoelastic properties of the material are represented by an equivalent circuit including a Newtonian damper and a linear elastic spring connected in parallel [Figure S2(a)], we have:

$$\frac{\partial x_{\rm s}(t)}{\partial t} = \frac{1}{\xi_{\rm s}}\left[F_{\rm c}(t) - k_{\rm s} x_{\rm s}(t)\right] \tag{A9}$$

Here $\xi_{\rm s}$ and $k_{\rm s}$ are constants that describe the viscous and elastic properties of the substrate.

Combining together Eq. (A7)-Eq. (A9) and Eq. (2), Eq. (6) and Eq. (7) from the main text, we finally obtain the following closed system of equations describing the dynamics of molecular clutches:

$$\begin{cases} \dfrac{\partial p_{\rm on}(l,t)}{\partial t} = -k_{\rm off}(l)\, p_{\rm on}(l,t) + k_{\rm on}(l)\, p_{\rm off}(t) - \left[v(t) - \dfrac{\partial x_{\rm s}(t)}{\partial t}\right] \dfrac{\partial p_{\rm on}(l,t)}{\partial l} \\ \dfrac{\partial p_{\rm off}(t)}{\partial t} = \displaystyle\int_{-\infty}^{+\infty} \left[k_{\rm off}(l)\, p_{\rm on}(l,t) - k_{\rm on}(l)\, p_{\rm off}(t)\right] {\rm d}l \\ \dfrac{\partial x_{\rm s}(t)}{\partial t} = \dfrac{1}{\xi_{\rm s}}\left[F_{\rm c}(t) - k_{\rm s} x_{\rm s}(t)\right] \\ p_{\rm off}(t) + \displaystyle\int_{-\infty}^{+\infty} p_{\rm on}(l,t)\, {\rm d}l = 1 \\ F_{\rm c}(t) = N_{\rm c} \displaystyle\int_{-\infty}^{+\infty} F(l)\, p_{\rm on}(l,t)\, {\rm d}l \\ v(t) = v_0 \left[1 - \dfrac{F_{\rm c}(t)}{N_{\rm m} F_{\rm st}}\right] \end{cases} \tag{A10}$$

The stationary solution of the above equation can be found by setting the left-hand sides of the first, second and third lines of the above equation equal to zero:

$$\begin{cases} \dfrac{\partial p_{\rm on}(l)}{\partial l} = -\dfrac{1}{v} k_{\rm off}(l)\, p_{\rm on}(l) + \dfrac{1}{v} k_{\rm on}(l)\, p_{\rm off} \\ \displaystyle\int_{-\infty}^{+\infty} k_{\rm off}(l)\, p_{\rm on}(l)\, {\rm d}l = \int_{-\infty}^{+\infty} k_{\rm on}(l)\, p_{\rm off}\, {\rm d}l \\ x_{\rm s} = \dfrac{F_{\rm c}}{k_{\rm s}} \\ p_{\rm off} + \displaystyle\int_{-\infty}^{+\infty} p_{\rm on}(l)\, {\rm d}l = 1 \\ F_{\rm c} = N_{\rm c} \displaystyle\int_{-\infty}^{+\infty} F(l)\, p_{\rm on}(l)\, {\rm d}l \\ v = v_0 \left[1 - \dfrac{F_{\rm c}}{N_{\rm m} F_{\rm st}}\right] \end{cases} \tag{A11}$$

Taking into account Eq. (3) from the main text, it is possible to further simplify Eq. (A11). First, it is clear that:

$$\int_{-\infty}^{+\infty} k_{\rm on}(l)\, p_{\rm off}\, {\rm d}l = k^0_{\rm on}\, p_{\rm off} \int_{-\infty}^{+\infty} \delta(l)\, {\rm d}l = k^0_{\rm on}\, p_{\rm off} \tag{A12}$$

Furthermore, by applying $\int_{-\infty}^{l'} {\rm d}l$ integration to the first line of Eq. (A10), where $l' < 0$, it can be found that:

$$\frac{\partial}{\partial t}\int_{-\infty}^{l'} p_{\rm on}(l,t)\, {\rm d}l = -\int_{-\infty}^{l'} k_{\rm off}(l)\, p_{\rm on}(l,t)\, {\rm d}l - \left[v(t) - \frac{\partial x_{\rm s}(t)}{\partial t}\right] p_{\rm on}(l',t), \ \text{if } l' < 0 \tag{A13}$$

Where we used the fact that $p_{\rm on}(-\infty, t) = 0$.

In the above equation, $k_{\rm off}(l)$, $p_{\rm on}(l,t)$ and $p_{\rm on}(l',t)$ are positive functions. In addition, model calculations show that near the stationary solutions given by Eq. (A11), we have: $v(t) - \frac{\partial x_{\rm s}(t)}{\partial t} \approx v(t) > 0$, see, for example, Figure 7 in the main text. I.e., molecular clutches are stretched in the positive direction of the x-axis due to the force generated by myosin II motor proteins [Figure S2(a)]. As a result, it is clear that the entire right-hand side of Eq. (A13) is negative, suggesting that near the stationary solutions the cumulative distribution function, ${\rm Pr}[l \le l'] = \int_{-\infty}^{l'} p_{\rm on}(l,t)\, {\rm d}l$, decreases with time in the case of $l' < 0$ until it reaches zero in the stationary state. Thus, in the stationary state:

$$p_{\rm on}(l) = 0, \ \text{if } l < 0 \tag{A14}$$

By combining Eq. (A11), Eq. (A12) and Eq. (A14), one can finally obtain Eq. (8) from the main text, which describes the stationary solution of Eq. (A10). This is the main formula, which was used to fit all experimental data in our work, since numerous experimental studies, as well as our own model calculations presented in Figures 7 and S7, suggest that myosin II motor proteins and the entire ensemble of molecular clutches operate close to a steady-state most of the time.

Indeed, using numerous experimental approaches, it was previously shown that the retrograde movement of actin filaments in living cells occurs at an almost constant speed ($v(t) \approx$ const) [4–7]. Taking into account this experimental fact, it can be seen from the last line of Eq. (A10) that $v(t) \approx$ const implies that $F_{\rm c}(t) \approx$ const. Then from the third line it can be concluded that $x_{\rm s}(t) \approx$ const as soon as $t \gg \tau_{\rm s}$, where $\tau_{\rm s} = \xi_{\rm s}/k_{\rm s}$ is the characteristic relaxation time of large-scale deformations of the substrate. Thus, applying $\frac{1}{T}\int_{t_0}^{t_0+T} {\rm d}t$ time-averaging to the first two lines of Eq. (A10) over a sufficiently long period of time, $T$ ($T \gg \max[\tau_{\rm s}, \tau_k]$, where $\tau_k$ is the characteristic relaxation time of the system due to the formation and dissociation of molecular clutches), we get:

$$\begin{cases} -k_{\rm off}(l)\, p_{\rm on}(l) + k_{\rm on}(l)\, p_{\rm off} - v\dfrac{\partial p_{\rm on}(l)}{\partial l} = \dfrac{1}{T}\displaystyle\int_{t_0}^{t_0+T} \dfrac{\partial p_{\rm on}(l,t)}{\partial t}{\rm d}t = \dfrac{p_{\rm on}(l, T+t_0) - p_{\rm on}(l,t_0)}{T} \xrightarrow[T\to\infty]{} 0 \\ \displaystyle\int_{-\infty}^{+\infty} \Big[k_{\rm off}(l)\, p_{\rm on}(l) - k_{\rm on}(l)\, p_{\rm off}\Big]\, {\rm d}l = \dfrac{1}{T}\int_{t_0}^{t_0+T} \dfrac{\partial p_{\rm off}(t)}{\partial t}{\rm d}t = \dfrac{p_{\rm off}(T+t_0) - p_{\rm off}(t_0)}{T} \xrightarrow[T\to\infty]{} 0 \end{cases} \tag{A15}$$

Where $p_{\rm on}(l) = \langle p_{\rm on}(l,t)\rangle_T = \frac{1}{T}\int_{t_0}^{t_0+T} p_{\rm on}(l,t){\rm d}t$ is the long time-averaged probability distribution. Similarly, $p_{\rm off} = \langle p_{\rm off}(t)\rangle_T = \frac{1}{T}\int_{t_0}^{t_0+T} p_{\rm off}(t){\rm d}t$. $t_0$ is an arbitrary moment in time.

In other words, from Eq. (A15) it follows that the long-term behaviour of the molecular clutch system is described by the stationary solution of Eq. (A10), which can be obtained by solving Eq. (8) from the main text.

The only question that remains is: how large must the time interval $T$ be for Eq. (8) to accurately describe the behaviour of the molecular clutch system, i.e., what is the typical time scale of the system relaxation to the stationary solution? Numeric calculations show that under the physiological values of the model parameters (Table T1), the molecular clutch system reaches the stationary state in $\sim 10-20$ s – a time-scale, which is close to experimentally measured characteristic times of formation and disassembly of molecular clutches, see Table T4 and the main text for more details. As a result, it can be concluded that Eq. (8) is suitable for description of cell adhesion processes on physiologically relevant time-scales of the order of tens of seconds or more.

Finally, we would like to note that the results shown in Figures 7 and S7 also indicate that even in the case of slowly relaxing viscous substrates ($\xi_{\rm s} \ge 1000$ pN·s/nm), the system still reaches a quasi-stationary state in $\sim 10-20$ s, which can still be very accurately described by the first five lines of Eq. (8) from the main text. Although the last (sixth) line of Eq. (8), which depicts the large-scale deformation of the substrate, is not satisfied, this does not affect the remaining results due to a negligible contribution of the slowly varying $x_{\rm s}(t)$ function to the time evolution of $p_{\rm on}(l,t)$ probability distribution, see Eq. (A10). Thus, it can be concluded that in the case when the substrate is made of a highly viscous material, $T \gg \max[\tau_{\rm s}, \tau_k]$ condition can be relaxed to a simpler one: $T \gg \tau_{\rm k}$, where $\tau_{\rm k}$ is of the order of several seconds.

### Appendix B: Stationary probability distribution

To derive the formula for the stationary probability distribution, $p_{\rm on}(l)$, it should be noted that the general solution to the first line of Eq. (8) in the main text has the following form:

$$p_{\rm on}(l) = Ce^{-\frac{1}{v}\int_0^l k_{\rm off}(l')\,{\rm d}l'} \tag{B1}$$

Where $C$ is an arbitrary constant. The exact value of this constant can be found by substituting Eq. (B1) into the third line of Eq. (8) and using the normalization condition of the probability distribution [Eq. (2)]. By doing so, it can be found that:

$$C = \frac{k_{\rm on}^0}{\displaystyle\int_0^{+\infty}\Big[k_{\rm off}(l') + k_{\rm on}^0\Big]e^{-\frac{1}{v}\int_0^{l'} k_{\rm off}(l''){\rm d}l''}{\rm d}l'} \tag{B2}$$

The above formula can be further simplified by noting that:

$$\int_0^{+\infty} k_{\rm off}(l')e^{-\frac{1}{v}\int_0^{l'} k_{\rm off}(l'')\,{\rm d}l''}{\rm d}l' = -v\int_0^{+\infty}{\rm d}\Big(e^{-\frac{1}{v}\int_0^{l'} k_{\rm off}(l'')\,{\rm d}l''}\Big) = v \tag{B3}$$

Here we have taken into account that $\int_0^0 k_{\rm off}(l'')\,{\rm d}l'' = 0$ and $\int_0^{+\infty} k_{\rm off}(l'')\,{\rm d}l'' = +\infty$.

By combining Eq. (B1)-Eq. (B3), it is straightforward to obtain Eq. (9) from the main text for the stationary probability distribution, $p_{\rm on}(l)$.

In the special case of slip-bonds ($k_{\rm off}^{0'} = 0$), which have a linear force-response given by Eq. (5), one can further streamline Eq. (9) as follows. First, from Eq. (4) and Eq. (5) it is clear that:

$$\int_0^l k_{\rm off}(l')\,{\rm d}l' = \int_0^l k_{\rm off}^0 e^{\beta k l' x_{\rm t}}\,{\rm d}l' = \frac{k_{\rm off}^0}{\beta k x_{\rm t}}\Big[e^{\beta k l x_{\rm t}} - 1\Big] \tag{B4}$$

Then, using the above formula, we have:

$$\int_0^{+\infty} e^{-\frac{1}{v}\int_0^l k_{\rm off}(l'){\rm d}l'}{\rm d}l = \left[a = \frac{k_{\rm off}^0}{\beta k v x_{\rm t}} \text{ and } \eta(l) = ae^{\beta k l x_{\rm t}}\right] = e^a\int_0^{+\infty} e^{-\eta(l)}{\rm d}l = \Big[{\rm d}\eta = \beta k x_{\rm t}\eta(l)\,{\rm d}l\Big] =$$

$$= \frac{e^a}{\beta k x_{\rm t}}\int_a^{+\infty}\eta^{-1}e^{-\eta}\,{\rm d}\eta = \frac{e^a}{\beta k x_{\rm t}}E_1(a) \tag{B5}$$

Where $E_1(a) = \int_a^{+\infty}\eta^{-1}e^{-\eta}{\rm d}\eta$ is the exponential integral function.

Finally, substituting Eq. (B4) and Eq. (B5) into Eq. (9) from the main text, it is easy to obtain Eq. (11) by showing that:

$$C = \frac{k_{\rm on}^0/v}{1 + \phi(a,\varepsilon)} \tag{B6}$$

Where $\phi(a,\varepsilon) = ae^{\varepsilon + a}E_1(a)$ and $\varepsilon = \ln(k_{\rm on}^0/k_{\rm off}^0)$.

Furthermore, using the same change of variables as in Eq. (B5), the following formula for the cell traction force ($F_{\rm c}$) can be obtained based on Eq. (5) and the fourth line of Eq. (8) from the main text:

$$F_{\rm c} = N_{\rm c}\int_0^{+\infty} F(l)\,p_{\rm on}(l)\,{\rm d}l = N_{\rm c}Ck\int_0^{+\infty} l\,e^{-\frac{1}{v}\int_0^l k_{\rm off}(l'){\rm d}l'}{\rm d}l = N_{\rm c}Cke^a\int_0^{+\infty} l\,e^{-\eta(l)}{\rm d}l =$$

$$= \frac{N_{\rm c}}{\beta x_{\rm t}}Ce^a\frac{\partial}{\partial s}\left(a^{-s}\int_0^{+\infty}\eta^s e^{-\eta(l)}{\rm d}l\right)\bigg|_{s=0} = \frac{N_{\rm c}}{k(\beta x_{\rm t})^2}Ce^a\frac{\partial}{\partial s}\Big(a^{-s}\Gamma(s,a)\Big)\bigg|_{s=0} \tag{B7}$$

Here $\Gamma(s,a) = \int_a^{+\infty}\eta^{s-1}e^{-\eta}{\rm d}\eta$ is the upper incomplete gamma function.

At the last stage, one just needs to use the formulas for the partial derivatives of $\Gamma(s,a)$ and $a^{-s}$ functions:

$$\frac{\partial\Gamma(s,a)}{\partial s} = \Gamma(s,a)\ln a + a\,G_{2,3}^{3,0}\left(\left.{0,\,0 \atop s-1,-1,-1}\right|\,a\right) \quad\text{and}\quad \frac{\partial a^{-s}}{\partial s} = -a^{-s}\ln a, \tag{B8}$$

Where $G_{p,q}^{m,n}$ is the Meijer's G-function, see Figure S3.

Then, taking into account that $\Gamma(0,a) = E_1(a)$, and substituting Eq. (B6) and Eq. (B8) into Eq. (B7), one can finally obtain Eq. (12) from the main text.

#### Appendix C: Talin clutch model

To account for the experimentally observed nonlinear behaviour of talin molecules and the force-induced unfolding of the mechanosensitive talin domain (R3), the first and the third line in Eq. (8) from the main text must be replaced by the following matrix equations:

$$\begin{cases} \dfrac{\partial \mathbf{p}_{\rm on}(l)}{\partial l} = \dfrac{1}{v}\mathbf{A}(l)\mathbf{p}_{\rm on}(l), \text{ for } l > 0 \\ \displaystyle\int_0^{+\infty} \mathbf{k}_{\rm off}^{\rm T}(l)\, \mathbf{p}_{\rm on}(l)\, {\rm d}l = p_{\rm off}\, k_{\rm on}^0 \end{cases} \tag{C1}$$

Where

$$\mathbf{p}_{\rm on}(l) = \begin{pmatrix} p_{\rm on}^{\rm f}(l) \\ p_{\rm on}^{\rm u}(l) \end{pmatrix} \quad \text{and} \quad \mathbf{k}_{\rm off}(l) = \begin{pmatrix} k_{\rm off}^{\rm f}(l) \\ k_{\rm off}^{\rm u}(l) \end{pmatrix} \quad \text{and} \quad \mathbf{A}(l) = \begin{pmatrix} -k_{\rm off}^{\rm f}(l) - k_{\rm u}(l) & k_{\rm f}(l) \\ k_{\rm u}(l) & -k_{\rm off}^{\rm u}(l) - k_{\rm f}(l) \end{pmatrix} \tag{C2}$$

Here $p_{\rm on}^{\rm f}(l)$ and $p_{\rm on}^{\rm u}(l)$ are the time-averaged distribution probabilities of molecular clutches in the case of folded and unfolded mechanosensitive talin domain, respectively. $k_{\rm off}^{\rm f}(l) = k_{\rm off}(F^{\rm f}(l))$ and $k_{\rm off}^{\rm u}(l) = k_{\rm off}(F^{\rm u}(l))$ are dissociation rates of molecular clutches in the case of folded and unfolded mechanosensitive talin domain, where $F^{\rm f}(l)$ and $F^{\rm u}(l)$ are the force-extension curves of molecular clutches in the corresponding states. $\mathbf{A}$ is the transition matrix. $k_{\rm u}(l) = k_{\rm u}(F^{\rm f}(l))$ and $k_{\rm f} = k_{\rm f}(F^{\rm u}(l))$ are the unfolding and refolding rates of the mechanosensitive talin domain. Superscript T denotes the matrix transpose.

In the model, $k_{\rm u}$ rate was approximated by the Bell's formula:

$$k_{\rm u}(l) = k_{\rm u}^0 e^{F^{\rm f}(l)/F_{\rm u}} \tag{C3}$$

Where $k_{\rm u}^0$ is the unfolding rate of the mechanosensitive talin domain at zero load. $F_{\rm u}$ is the characteristic unfolding force of the mechanosensitive talin domain.

As for $k_{\rm f}$, it was calculated from the equilibrium constant ($K_{\rm eq}$):

$$K_{\rm eq}(l) = \frac{k_{\rm f}(l)}{k_{\rm u}(l)} = e^{-\Delta G(l)/k_{\rm B}T} \tag{C4}$$

Where $\Delta G$ is the difference between the free energies of a molecular clutch in the cases of the folded and unfolded mechanosensitive talin domain:

$$\Delta G(l) = -k_{\rm B}T \ln \frac{k_{\rm f}^0}{k_{\rm u}^0} + \int_0^l \Big[F^{\rm f}(l') - F^{\rm u}(l')\Big] {\rm d}l' \tag{C5}$$

Here $k_{\rm f}^0$ is the refolding rate of the mechanosensitive talin domain at zero load.

To calculate the force-extension curves $F^{\rm f}(l)$ and $F^{\rm u}(l)$, we used the freely-jointed chain model for the rod part of talin, treating each globular domain as a solid body. As for the talin unstructured peptide linker and unfolded mechanosensitive talin domain, they were described with the help of the worm-like chain model.

In order to solve Eq. (C1), we first applied the Runge-Kutta method to the differential equation from the first line, which describes the evolution of the vector $\mathbf{p}_{\rm on}$ as a function of the molecular clutch extension, $l$. To this aim, we used the following boundary condition:

$$\widetilde{\mathbf{p}}_{\rm on}(0) = \frac{1}{k_{\rm f}(0) + k_{\rm u}(0)} \begin{pmatrix} k_{\rm f}(0) \\ k_{\rm u}(0) \end{pmatrix} \tag{C6}$$

The resulting solution, $\widetilde{\mathbf{p}}_{\rm on}(l)$, was then renormalized to obtain the final probability distribution, $\mathbf{p}_{\rm on}(l)$, satisfying the first and the second lines of Eq. (C1):

$$\mathbf{p}_{\rm on}(l) = \frac{k_{\rm on}^0 \widetilde{\mathbf{p}}_{\rm on}(l)}{\displaystyle\int_0^{+\infty} \Big[\mathbf{k}_{\rm off}^{\rm T}(l') + (k_{\rm on}^0, k_{\rm on}^0)\Big] \widetilde{\mathbf{p}}_{\rm on}(l')\, {\rm d}l'} = \frac{k_{\rm on}^0 \widetilde{\mathbf{p}}_{\rm on}(l)}{v + \displaystyle\int_0^{+\infty} (k_{\rm on}^0, k_{\rm on}^0) \cdot \widetilde{\mathbf{p}}_{\rm on}(l')\, {\rm d}l'} \tag{C7}$$

Finally, the traction force generated by engaged molecular clutches was calculated as:

$$F_{\rm c} = \frac{N_{\rm c} \displaystyle\int_0^{+\infty} \mathbf{F}^{\rm T}(l)\, \mathbf{p}_{\rm on}(l)\, {\rm d}l}{1 - \alpha \displaystyle\int_0^{+\infty} p_{\rm on}^{\rm u}(l')\, {\rm d}l'} \tag{C8}$$

Where $\mathbf{F}^{\mathrm{T}}(l) = (F^{\mathrm{f}}(l), F^{\mathrm{u}}(l))$, and $\alpha$ is the model parameter describing the force-induced reinforcement of cell adhesion sites [8] due to exposure of the talin's cryptic site for vinculin binding [9, 10]. Specifically, $\alpha$ is the average number of new sites available for the molecular clutch formation, which are added to the system upon unfolding of each mechanosensitive talin domain.

It should be noted that although in this study we considered the force-induced unfolding of only one mechanosensitive talin domain (R3), the developed theoretical framework allows full-scale simulation by considering the entire network of folding-unfolding transitions of all talin domains.

#### Appendix D: Linear correlation between cell traction and retrograde actin flow

From Figures 3(a,b,d,e) in the main text it appears that all retrograde actin flow curves reflected in the vertical direction are very similar in shape to the corresponding cell traction curves. Indeed, by plotting cell traction as a function of retrograde actin flow, it was found that there is a negative linear correlation between the two quantities, which is confirmed by experimental data, see Figure 3(f). This interesting phenomenon originates from the linear force-velocity relationship of myosin II motors, described by Eq. (7) in the main text. Indeed, the cell traction and retrograde actin flow curves are obtained in the model by projecting the intersection of two surfaces corresponding to the cell traction force ($F_{\mathrm{c}}$) and myosin II-pulling force ($F_{\mathrm{m}}$) in vertical and horizontal directions, respectively, see Figure 2(c). Since on a linear scale the surface corresponding to the myosin II-pulling force has the shape of a plane described by the linear force-velocity relationship [Eq. (7)], it is clear that the projection of the intersection curve belonging to this plane in the horizontal and vertical directions will result in a negative linear correlation between the cell traction and retrograde actin flow curves. Thus, the experimental observation of such a correlation indicates that myosin II motors do indeed have a linear force-velocity relationship in living cells.

#### Appendix E: Average tension of molecular clutches vs the area of cell adhesion

Previously, using a FRET talin tension sensor, it was found in experiments that the average tension experienced by talin molecules in FAs formed by mouse embryonic stem cells does not depend on either the total amount of talin in FAs or their size [11]. Although the molecular origin of this behaviour of FAs is still unclear, the linear and talin clutch models developed in our study can be used to potentially understand this phenomenon.

Indeed, from Figure S4(d) it is clear that even if the rate of formation of talin-based molecular clutches in the WT-2 model is decreased by 10 times, which corresponds to a 10 times smaller density of talins in FA, this leads to an increase in the tension of most talin-based molecular clutches by $\leq 50\%$. This result suggests that at the initial stage of FA maturation talin molecules in molecular clutches experience a fairly constant tension, which is in good agreement with experimental data. As for more mature FAs that have already reached the maximum density of talins (i.e., $> 2$ min after the start of their formation [12]), an even stronger and more general result can be proven in a mathematically rigorous way.

Specifically, it should first be noted that from the fourth line of Eq. (8) it follows that the average tension experienced by each molecular clutch ($F_1$) does not depended explicitly on the total number of binding sites ($N_{\mathrm{c}}$) available for the formation of molecular clutches in the cell adhesion area:

$$F_1 = \frac{F_{\mathrm{c}}}{N_{\mathrm{e}}} = \frac{\displaystyle\int_0^{+\infty} F(l)\, p_{\mathrm{on}}(l)\, \mathrm{d}l}{\displaystyle\int_0^{+\infty} p_{\mathrm{on}}(l)\, \mathrm{d}l} \tag{E1}$$

Here $N_{\mathrm{e}} = N_{\mathrm{c}} \int_0^{+\infty} p_{\mathrm{on}}(l)\, \mathrm{d}l$ is the average total number of molecular clutches in the cell adhesion area.

On the other hand, it follows from Eq. (9) that the only way in which $p_{\mathrm{on}}(l)$ probability distribution of molecular clutch extension ($l$) may potentially depend on the cell adhesion area is through the rate of the retrograde actin flow, $v$ [i.e., $p_{\mathrm{on}}(l) = p_{\mathrm{on}}(l, v)$]. In turn, the rate of the retrograde actin flow is determined by the following implicit formula, which can be derived from the fourth and fifth lines of Eq. (8):

$$\frac{N_{\mathrm{c}}}{N_{\mathrm{m}}} \int_0^{+\infty} F(l)\, p_{\mathrm{on}}(l, v)\, \mathrm{d}l = F_{\mathrm{st}} \left[1 - \frac{v}{v_0}\right] \tag{E2}$$

The only variables that depend on the cell adhesion area ($A = \pi R^2$) in the above formula are $N_{\mathrm{m}}$ and $N_{\mathrm{c}}$: $N_{\mathrm{m}} = A\sigma_{\mathrm{m}}$ and $N_{\mathrm{c}} = A\sigma_{\mathrm{c}}$, where $R$ is the characteristic radius of the cell adhesion area, and $\sigma_{\mathrm{m}}$ and $\sigma_{\mathrm{c}}$ are the densities of myosin II motors and sites available for the formation of molecular clutches in the cell adhesion area, respectively. Yet, it is obvious that $N_{\rm c}/N_{\rm m}$ ratio does not depend on the cell adhesion area as soon as $\sigma_{\rm m}$ and $\sigma_{\rm c}$ are constant, since both $N_{\rm m}$ and $N_{\rm c}$ are proportional to the cell adhesion area. Thus, from Eq. (E1)-Eq. (E2) and Eq. (9) it can be concluded that neither the rate of the retrograde actin flow ($v$), nor the stationary probability distribution of molecular clutch extension ($p_{\rm on}$), nor the average tension experienced by each molecular clutch ($F_1$) depend on the cell adhesion area, which is in good agreement with previous experimental observations [11].

Finally, it should be noted that although in this Appendix section we relied on formulas describing linear molecular clutches, the same reasoning can be used to reach a similar conclusion within the framework of the talin WT model.

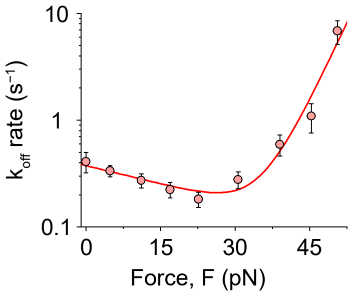


FIG. S1. **Integrin dissociation rate from fibronectin as a function of the applied mechanical load.** Solid curve shows two-term exponential fitting [Eq. (4) in the main text] of experimental data points taken from ref. [6].

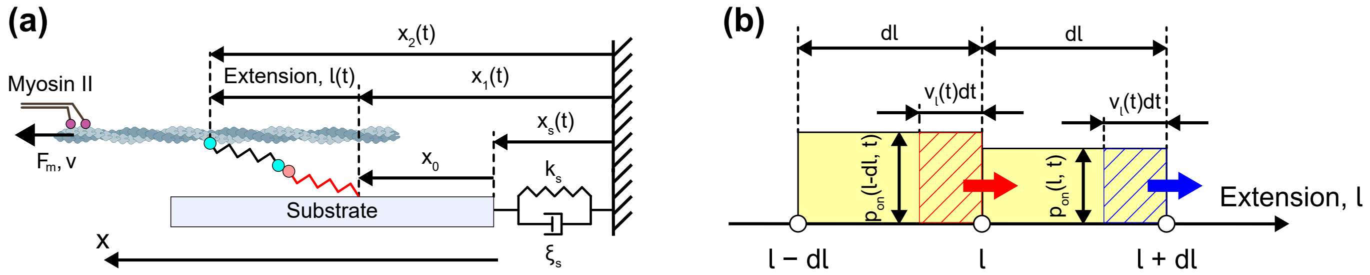


FIG. S2. **Molecular clutch extension and its probability distribution.** **(a)** In the model, extension of molecular clutches is defined as the distance between their both ends along the x-axis: $l(t) = x_2(t) - x_1(t)$, where $x_1(t)$ and $x_2(t)$ are the x-coordinates of the ends of a molecular clutch. In turn, $x_1(t)$ and $x_2(t)$ coordinates can be expressed in terms of the speed of the retrograde actin flow, $v(t)$, the substrate deformation, $x_s(t)$, and the coordinate of the molecular clutch attachment point to the substrate, $x_0$, as discussed in Appendix A. **(b)** To derive the master equation describing the time evolution of the probability distribution of molecular clutch extension, $p_{\rm on}(l,t)$, one needs to consider the axis showing all possible extensions of molecular clutches, dividing it into infinitesimal intervals of d$l$ length, such that the probability distribution $p_{\rm on}(l,t)$ can be considered uniform on each of these intervals, as schematically shown in the figure. Then the probability to observe a molecular clutch at a given site with an extension in $[l, l + {\rm d}l]$ interval is equal to $p_{\rm on}(l,t)\,{\rm d}l$, which is simply the area of the right yellow rectangle shown in the figure. Over a small time step, d$t$, this probability changes due to the gradual stretching of molecular clutches at $v_l(t)$ rate caused by the retrograde flow of actin filaments and substrate deformation, see Eq. (A1)-Eq. (A3). The net change in the probability because of the influx of newly stretched molecular clutches from the left side of $[l, l + {\rm d}l]$ interval (red striped box) and outflow from the right side (blue striped box) is ${\rm d}p_{\rm on}(l,t)|_{\rm stretch} \cdot {\rm d}l = [p_{\rm on}(l - {\rm d}l, t) - p_{\rm on}(l,t)]\, v_l(t)\, {\rm d}t$.

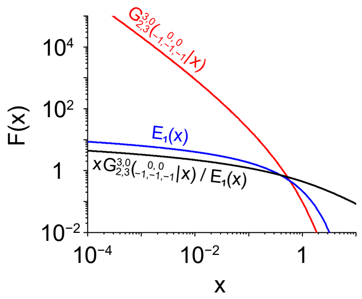


FIG. S3. **Graphs of the Meijer's $G^{3,0}_{2,3}$ function and the exponential integral function, $E_1$.** The figure shows graphs of $G^{3,0}_{2,3}\left(\begin{smallmatrix}0,\ 0\\ -1,-1,-1\end{smallmatrix}\middle|\, x\right)$ and $E_1(x)$ functions, as well as $xG^{3,0}_{2,3}\left(\begin{smallmatrix}0,\ 0\\ -1,-1,-1\end{smallmatrix}\middle|\, x\right)/E_1(x)$ term from Eq. (12) in the main text. $G^{3,0}_{2,3}\left(\begin{smallmatrix}0,\ 0\\ -1,-1,-1\end{smallmatrix}\middle|\, x\right)$ and $E_1(x)$ were calculated using built-in Matlab functions, meijerG([ ], $[0, 0]$, $[-1, -1, -1]$, [ ], $x$) and expint($x$), respectively.

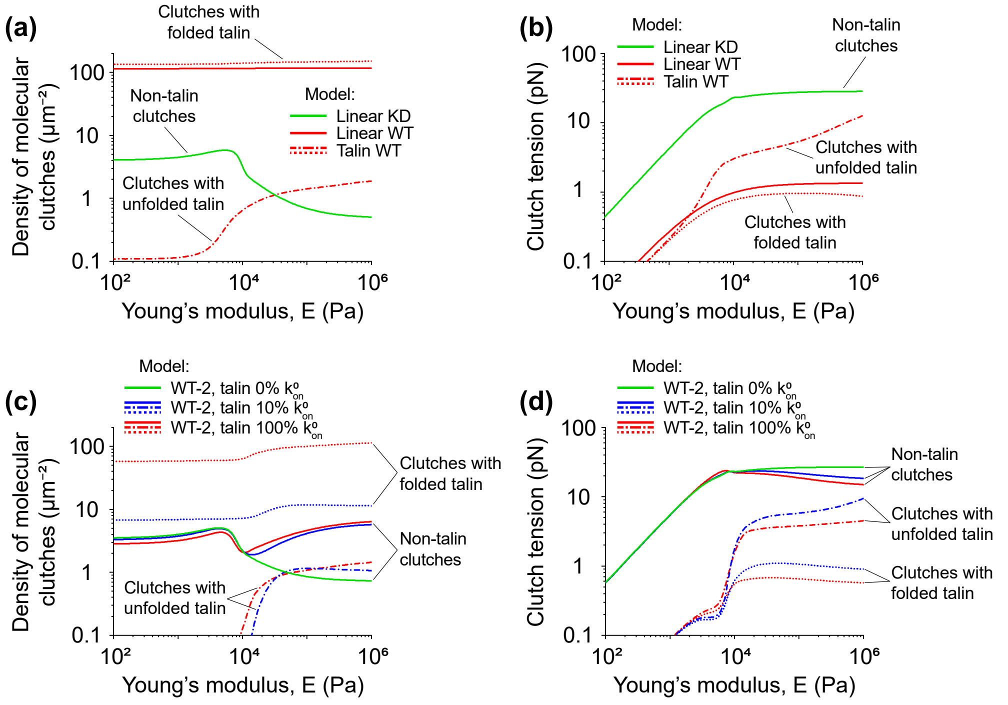


FIG. S4. **Average density and tension of molecular clutches. (a-d)** Mean densities (a, c) and average tensions (b, d) of molecular clutches predicted by the different models tested in our study. In panels (a-d), the dotted and dash-dotted curves indicate talin-based molecular clutches with the folded and unfolded talin mechanosensitive R3 domain, respectively. Green and red solid curves in panels (a, b) represent non-talin-based and talin-based molecular clutches in the linear KD and linear WT models, respectively; whereas the solid curves in panels (c, d) correspond to non-talin-based molecular clutches in the WT-2 model. From panels (a, c) it can be seen that only $1\% - 2\%$ of talin-based molecular clutches experience forces high enough to induce unfolding of the mechanosensitive R3 domain of talin, in good agreement with experimental studies performed on *Xenopus laevis* cells grown on a fibronectin-coated glass substrate, in which only about $\sim 4\%$ of talin molecules exhibited significant conformational changes upon molecular clutch formation [13]. Furthermore, the figure shows that the average tension experienced by talin-based molecular clutches increases with substrate rigidity up to a value of the substrate Young's modulus of $\sim 10$ kPa, after which it reaches a plateau in $0.5 - 1.3$ pN and $3 - 12$ pN force range in the case of molecular clutches with the folded and unfolded mechanosensitive R3 domain of talin, respectively. This model prediction agrees well with previous *in vivo* experimental studies based on the FRET talin tension sensor [11, 14, 15]. The results shown in panels (a, b) and panels (c, d) were obtained using the model parameter values listed in Table T1 and Table T2, respectively.

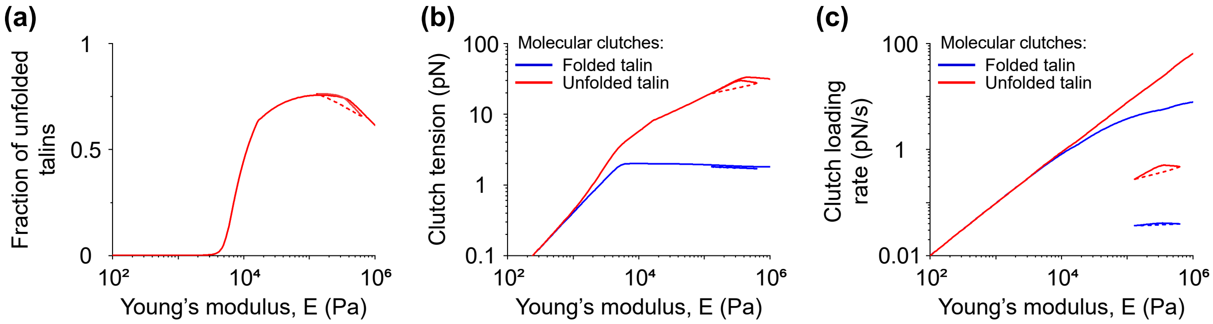


FIG. S5. **Fraction of unfolded talins, average tension and loading rate of molecular clutches in the talin WT model with a reduced dissociation rate of molecular clutches and a lower density of sites available for the formation of molecular clutches. (a-c)** Fraction of unfolded talins (a), average tensions (b) and loading rates (c) of talin-based molecular clutches as a function of the Young's modulus of the substrate ($E$). In the plots, solid curves indicate stable stationary branches of the graphs, and dashed curves show unstable branches. In panels (b, c), red curves represent molecular clutches with the unfolded mechanosensitive R3 domain of talin, whereas blue curves indicate molecular clutches with folded talin. The results shown in panels (a-c) were obtained using the talin WT model parameter values listed in Table T1, with two exceptions – the values of $k_{\text{off}}^{0}$ and $k_{\text{off}}^{0'}$ parameters were 100 times less than those shown in the table, and $\sigma_{\text{c}}$ was set to 40 $\mu$m$^{-2}$. These values were chosen to reflect the typical range of integrin dissociation rates measured in experiments [16, 17] and the wide range of densities of sites available for molecular clutch formation that are often used in mechanosensitive cell adhesion experiments [18, 19]. The graphs show that under such conditions the fraction of unfolded talin molecules increases to $\sim 70\%$ on rigid substrates ($E > 5$ kPa), and the average tension and loading rate experienced by molecular clutches becomes as high as 30 pN and 60 pN/s, respectively, suggesting that the dissociation rate of molecular clutches and their density can strongly influence these parameters.

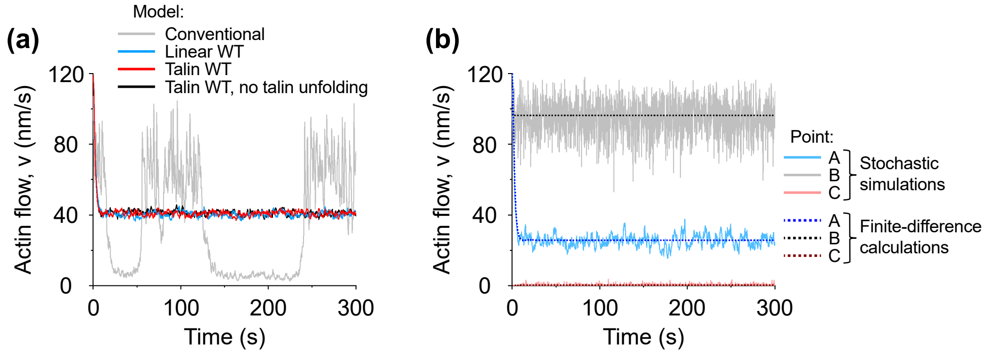


FIG. S6. **Stochastic simulations and finite-difference calculations of the time evolution of the molecular clutch system.** **(a)** Representative trajectories predicted by the conventional molecular clutch model (gray color), the linear WT model (blue color), the talin WT model (red color), and the talin WT model in the absence of the force-induced unfolding of the mechanosensitive talin domain (black color), obtained using stochastic simulations. Calculations were carried out for 10 kPa Young's modulus of the substrate. As can be seen from the graph, the conventional molecular clutch model predicts strong fluctuations in retrograde actin flow under such conditions, whereas the linear WT and talin WT models demonstrate a much milder behaviour of the molecular clutch system, which is in better agreement with experimental observations showing that retrograde actin flow does not experience strong fluctuations in cells [4–7]. In addition, the graph shows that the force-induced unfolding of the mechanosensitive R3 domain of talin does not have a strong effect either on the relaxation dynamics of the molecular clutch system to the stationary state, or on the stationary value of retrograde actin flow. The values of the model parameters used in the calculations of the linear WT and talin WT models are given in Table T1. As for the conventional molecular clutch model, the values of the main model parameters were taken from ref. [6], except of the spring constant of the molecular clutches, which was set to 1 pN/nm. **(b)** Representative trajectories demonstrating the time evolution of the molecular clutch system obtained by solving the master equation [Eq. (A10)] for points A, B and C shown in Figure 4(b) in the main text by using stochastic simulations and finite-difference calculations. From the graph and Figure 4(b) it can be seen that if at small values of the Young's modulus of the substrate ($E \lesssim 2$ MPa) there is a unique stationary solution that attracts trajectories to its neighbourhood (point A), then at higher values of Young's modulus ($E \gtrsim 2$ MPa) there is a pair of such stationary solutions (points B and C), which indicates the bistable behaviour of the molecular clutch system. Notably, stochastic simulations indicate that both strong and weak cell adhesion states are very stable and can last for many minutes without experiencing stochastic transition between themselves.

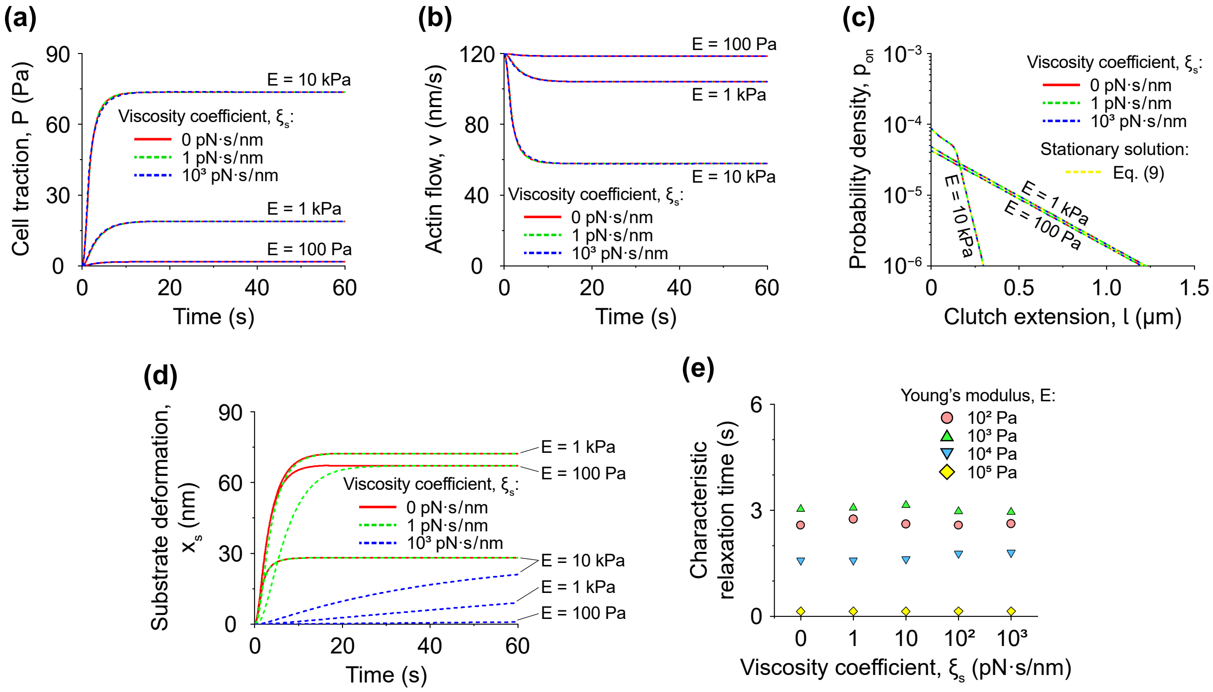


FIG. S7. **Time evolution of cell adhesion in MEF Talin 1 KO, Talin 2 shRNA cells.** **(a, b)** Time evolution of cell traction ($P$), retrograde actin flow ($v$) and substrate deformation ($x_\mathrm{s}$), starting from an initial configuration with zero molecular clutches between cell and substrate. The curves were obtained by numerically solving the master equation describing the time evolution of the system [Eq. (A10)] using the finite-difference method. Calculations were carried out for three different values of the substrate Young's modulus ($E$): 100 Pa, 1 kPa and 10 kPa. The substrate viscosity coefficient ($\xi_\mathrm{s}$) in the same calculations was varied from 0 pN·s/nm to 1000 pN·s/nm. The values of the remaining model parameters were the same as those in the linear KD model, see Table T1. It can be seen that cell traction and retrograde actin flow quickly reach stationary values within $\sim 10-20$ s, regardless of the substrate viscosity. **(c)** Probability distribution ($p_\mathrm{on}$) of molecular clutch extension ($l$), calculated after 20 s time evolution of the molecular clutch system for three different values of Young's modulus ($E$) and viscosity coefficient of the substrate ($\xi_\mathrm{s}$). The graph shows that the molecular clutch system reaches the stationary / quasi-stationary probability distribution described by Eq. (9) from the main text (yellow dashed lines) within 20 s, regardless of the substrate viscosity. **(d)** Time evolution of substrate deformation. As can be seen from the graph, the characteristic relaxation time of the substrate is highly dependent on its viscosity, ranging from a few seconds to many minutes. Yet, from panels (a-c) it is clear that slow substrate relaxation has little effect on the rapid convergence of the probability distribution of molecular clutch extension, as well as cell traction and actin retrograde flow to their stationary / quasi-stationary values / states. **(e)** Dependence of the characteristic relaxation time of retrograde actin flow on Young's modulus and viscosity coefficient of the substrate. The characteristic relaxation time was calculated by fitting the curves shown in panel (b) to an exponential decay function. As can be seen from the graph, the characteristic relaxation time decreases with increasing Young's modulus of the substrate. However, it is practically independent of the viscous properties of the substrate.

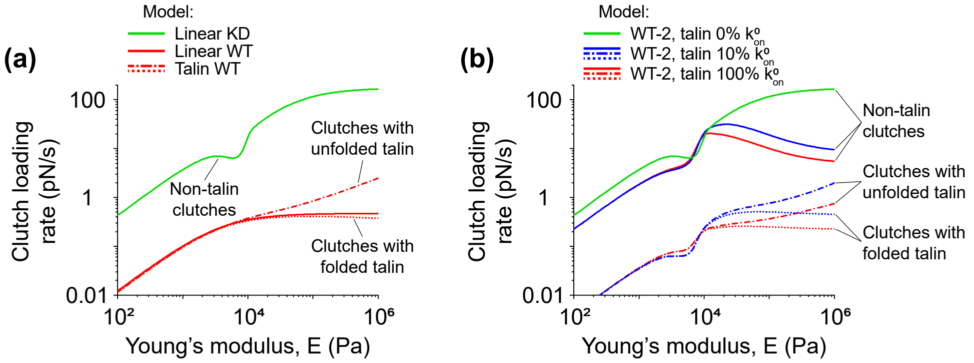


FIG. S8. **Average loading rate of molecular clutches. (a, b)** Average loading rate of molecular clutches predicted by the linear KD, linear WT, talin WT and WT-2 models. In panels (a, b), the dotted and dash-dotted curves indicate talin-based molecular clutches with the folded and unfolded talin mechanosensitive R3 domain, respectively. Green and red solid curves in panel (a) represent non-talin-based and talin-based molecular clutches in the linear KD and linear WT models, respectively; whereas the solid curves in panel (b) correspond to non-talin-based molecular clutches in the WT-2 model. The figure shows that the average loading rate of molecular clutches in MEF WT cells predicted by the linear WT, talin WT and WT-2 models can reach $0.2 - 5.5$ pN/s on rigid substrates ($E = 1$ MPa), in good agreement with the loading rate of $0.5 - 4$ pN/s measured in a recent experimental study [19]. On the other hand, the average loading rate of molecular clutches in MEF Talin 1 KO, Talin 2 shRNA cells predicted by the linear KD model can be as high as 170 pN/s [green curve in panel (a)]. Interestingly, calculations from the WT-2 model show that even 10% of the maximum density of talin in cell adhesion areas and therefore 10% of the rate of talin-based molecular clutch formation ($k^0_{\mathrm{on}}$) is sufficient to greatly reduce the loading rate of molecular clutches [compare the green and blue solid curves in panel (b)], which indicates a very strong effect of talin molecules on stabilizing cell adhesion. The results shown in panels (a, b) were obtained using the model parameter values listed in Table T1 and Table T2, respectively.

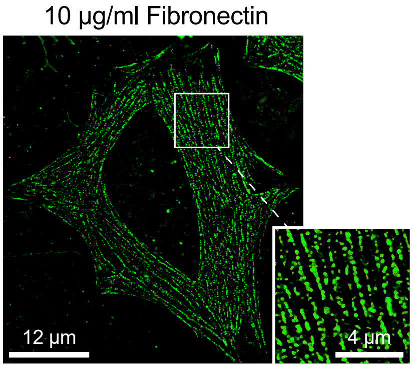


FIG. S9. **SIM image of HeLa cells expressing myosin RLC-GFP construct.** The inset shows a magnified area indicated by the white square in the figure. From the figure it can be seen that myosin II stacks form an almost uniform network in HeLa cells, resembling beads on strings.

TABLE T1. Parameter values in the linear KD, linear WT and talin WT models.

| Parameter | Model [1] | | | Description |
|---|---|---|---|---|
| | **Linear KD** | **Linear WT** | **Talin WT** | |
| $k^0_{\mathrm{on}}$ | 0.005 $\mathrm{s}^{-1}$ | 0.23 $\mathrm{s}^{-1}$ | 0.3 $\mathrm{s}^{-1}$ | Assembly rate of molecular clutches in a mechanically relaxed state (fitted). The estimated values are in fairly good agreement with the experimentally measured rates of formation of talin-based molecular clutches in *Xenopus laevis* cells (0.03 – 0.06 $\mathrm{s}^{-1}$ [13]). |
| $k^0_{\mathrm{off}}, k^{0\prime}_{\mathrm{off}}$ | —"— | 5.4·10$^{-5}$ $\mathrm{s}^{-1}$ and 0.38 $\mathrm{s}^{-1}$ [6] | —"— | Catch-bond dissociation rates of molecular clutches at zero mechanical load, see Figure S1. |
| $x_{\mathrm{t}}, x'_{\mathrm{t}}$ | —"— | 0.97 nm and −0.11 nm [6] | —"— | Catch-bond transition state distances of molecular clutches, see Figure S1. |
| $v_0$ | —"— | 120 nm/s [4, 6, 20, 21] | —"— | Movement rate of myosin II motor proteins in the absence of mechanical load. |
| $F_{\mathrm{st}}$ | —"— | 2 pN [22] | —"— | Stalling force of myosin II motor proteins. |
| $\sigma_{\mathrm{m}}$ | 71 $\mu\mathrm{m}^{-2}$ | 85 $\mu\mathrm{m}^{-2}$ | 82 $\mu\mathrm{m}^{-2}$ | Density of myosin II motor proteins near the cell-substrate interface (fitted). The estimated values are close to the experimentally measured density of myosin II filaments ($\sim 2-3$ $\mu\mathrm{m}^{-2}$ [7, 23]) multiplied by the average number of individual myosin molecules in each myosin II filament ($\sim$ 30 [24]). |
| $\sigma_{\mathrm{c}}$ | —"— | 300 $\mu\mathrm{m}^{-2}$ [6] | —"— | Density of sites available for the molecular clutch formation near the cell-substrate interface. |
| $k_{\mathrm{c}}$ | 1.6 pN/nm | 0.05 pN/nm | | Spring constant of the intracellular part of molecular clutches (fitted, linear KD and linear WT models only). Experimentally measured spring constant of talin in the physiologically relevant range of 0 – 10 pN is $\sim$ 0.1 pN/nm, see Table T3. |
| $r$ | 26 nm | 0.8 nm | 0.6 nm | Characteristic radius of the adhesion site of a single molecular clutch (fitted). |
| $R$ | —"— | 1700 nm [6] | —"— | Characteristic radius of cell adhesion areas. |
| $l_{\mathrm{y}}$ | 150 nm | $>$ 300 nm | | Molecular clutch extension corresponding to the yield strength (fitted, linear KD and linear WT models only). The value of this model parameter did not have a strong effect on the results in the case of the linear WT model. |
| $L$ | | | 37 nm [25] | Length of the talin unstructured peptide linker. |
| $L_{\mathrm{p}}$ | | | 1 nm [26] | Average persistence length of a polypeptide chain. |
| $N_{\mathrm{g}}$ | | | 12 [25] | Total number of the force-transmitting globular domains in the rod part of talin. |
| $b$ | | | 5 nm | Average size of the talin globular domains. |
| $k^0_{\mathrm{u}}$ | | | 0.018 $\mathrm{s}^{-1}$ [10] | Unfolding rate of the mechanosensitive talin domain (R3) at zero load. |
| $F_{\mathrm{u}}$ | | | 0.7 pN [10] | Characteristic unfolding force of the mechanosensitive talin domain. |
| $k^0_{\mathrm{f}}$ | | | 22.2 $\mathrm{s}^{-1}$ [10] | Refolding rate of the mechanosensitive talin domain at zero load. |
| $\alpha$ | | | 2 | Average number of new sites available for the formation of molecular clutches added to the system upon unfolding of a single mechanosensitive talin domain (fitted). The value of this model parameter did not have a strong effect on the results. |

[1] In the linear KD and linear WT models describing MEF Talin 1 KO, Talin 2 shRNA cells and MEF WT cells, respectively, molecular clutches were represented by two-part linear springs up to the yield point, $l_{\mathrm{y}}$, see Eq. (13) in the main text. In the talin WT model describing MEF WT cells, the intracellular parts of molecular clutches were represented by nonlinear springs with a talin-like force-response, whereas the extracellular part was represented by a linear spring, see Methods and Appendix C.

TABLE T2. Values of fitting parameters in the WT-2 model [1,2].

| Parameter | General | Molecular clutches | | Description |
|---|---|---|---|---|
| | | Non-talin-based | Talin-based | |
| $k_{\mathrm{on}}^{0}$ | | 0.004 $\mathrm{s}^{-1}$ | 0.091 $\mathrm{s}^{-1}$ | Assembly rate of molecular clutches in a mechanically relaxed state. |
| $\sigma_{\mathrm{m}}$ | 88 $\mu\mathrm{m}^{-2}$ | | | Density of myosin II motor proteins near the cell-substrate interface. |
| $k_{\mathrm{c}}$ | | 0.9 pN/nm | | Spring constant of the intracellular part of molecular clutches. |
| $r$ | | 33 nm | 0.3 nm | Characteristic radius of the adhesion site of a single molecular clutch. |
| $l_{\mathrm{y}}$ | | 160 nm | | Molecular clutch extension corresponding to the yield strength. |
| $\alpha$ | | | 200 | Average number of new sites available for the formation of molecular clutches added to the system upon unfolding of a single mechanosensitive talin domain. The value of this model parameter did not have a strong effect on the results. |

[1] In the WT-2 model describing MEF WT cells, the intracellular parts of talin-based molecular clutches were represented by nonlinear springs with a talin-like force-response [Appendix C], whereas the intracellular parts of non-talin-based molecular clutches were represented by linear springs (up to the yield point, $l_{\mathrm{y}}$) [see Methods section in the main text].

[2] The table shows only the values of the fitting parameters. The remaining model parameters had the same values as in the linear KD and talin WT models displayed in Table T1.

TABLE T3. Experimentally measured elasticity of cytockeletal and focal adhesion proteins in the force range of $0 - 10$ pN.

| **Protein constructs** | **Average elasticity** |
|---|---|
| Talin 1, rod domain | 0.12 pN/nm [10] |
| Talin 1, rod domain + talin unstructured linker | 0.09 pN/nm (estimated based on the WLC model) |
| Full-length vinculin | 0.12 pN/nm [27] |
| Full-length $\alpha$-actinin 1 | 0.19 pN/nm [28] |
| Filamin A, rod 1 part, IgFLNa 1-8 domains | 0.28 pN/nm [29] |

TABLE T4. Characteristic time scales of cell adhesion processes.

| **Process** | **Characteristic time** |
|---|---|
| Relaxation of the molecular clutch system to the stationary / quasi-stationary state | 0.1 – 4 s, this study |
| Formation and transition of talin-based molecular clutches to a stretched state | 18 – 32 s [13] |
| Dissociation of talin-based molecular clutches | 1.5 – 2.7 s [13] |
| Dissociation of integrin from fibronectin | 1 – 5 s [6] |
| Talin turnover in focal adhesions | 15 – 50 s [11, 14, 15, 30] |
| Integrin turnover in focal adhesions | 20 – 30 s [30] |
| Binding / dissociation of the minor (dynamic) fraction of paxillin and vinculin to / from focal adhesions | 30 – 60 s [31] |
| Cell spreading | 6 min [32] |
| Focal adhesions, average lifetime | 10 – 90 min [31, 33] |
| Dissociation of the major (stably bound) fraction of paxillin and vinculin from focal adhesions | $>$ 30 min [31] |

[1] M. Srinivasan and S. Walcott, Binding site models of friction due to the formation and rupture of bonds: state-function formalism, force-velocity relations, response to slip velocity transients, and slip stability, Phys. Rev. E **80**, 046124 (2009).
[2] B. Sabass and U. S. Schwarz, Modeling cytoskeletal flow over adhesion sites: competition between stochastic bond dynamics and intracellular relaxation, J. Phys.: Condens. Matter **22**, 194112 (2010).
[3] Y. Li, P. Bhimalapuram, and A. R. Dinner, Model for how retrograde actin flow regulates adhesion traction stresses, J. Phys.: Condens. Matter **22**, 194113 (2010).
[4] C. E. Chan and D. J. Odde, Traction dynamics of filopodia on compliant substrates, Science **322**, 1687 (2008).
[5] T. Bornschlögl, S. Romero, C. L. Vestergaard, J.-F. Joanny, G. Tran Van Nhieu, and P. Bassereau, Filopodial retraction force is generated by cortical actin dynamics and controlled by reversible tethering at the tip, Proc. Natl. Acad. Sci. U.S.A. **110**, 18928 (2013).
[6] A. Elosegui-Artola, R. Oria, Y. Chen, A. Kosmalska, C. Pérez-González, N. Castro, C. Zhu, X. Trepat, and P. Roca-Cusachs, Mechanical regulation of a molecular clutch defines force transmission and transduction in response to matrix rigidity, Nat. Cell Biol. **18**, 540 (2016).
[7] N. O. Alieva, A. Efremov, S. Hu, D. Oh, Z. Chen, M. Natarajan, H. T. Ong, A. Jégou, G. Romet-Lemonne, J. T. Groves, M. P. Sheetz, J. Yan, and A. D. Bershadsky, Myosin IIA and formin dependent mechanosensitivity of filopodia adhesion, Nat. Commun. **10**, 3593 (2019).
[8] D. Riveline, E. Zamir, N. Q. Balaban, U. S. Schwarz, T. Ishizaki, S. Narumiya, Z. Kam, B. Geiger, and A. D. Bershadsky, Focal contacts as mechanosensors: externally applied local mechanical force induces growth of focal contacts by an mDia1-dependent and ROCK-independent mechanism, J. Cell Biol. **153**, 1175 (2001).
[9] M. Yao, B. T. Goult, H. Chen, P. Cong, M. P. Sheetz, and J. Yan, Mechanical activation of vinculin binding to talin locks talin in an unfolded conformation, Sci. Rep. **4**, 4610 (2014).
[10] M. Yao, B. T. Goult, B. Klapholz, X. Hu, C. P. Toseland, Y. Guo, P. Cong, M. P. Sheetz, and J. Yan, The mechanical response of talin, Nat. Commun. **7**, 11966 (2016).
[11] A. Kumar, M. Ouyang, K. Van den Dries, E. J. McGhee, K. Tanaka, M. D. Anderson, A. Groisman, B. T. Goult, K. I. Anderson, and M. A. Schwartz, Talin tension sensor reveals novel features of focal adhesion force transmission and mechanosensitivity, J. Cell Biol. **213**, 371 (2016).
[12] A. I. Bachir, J. Zareno, K. Moissoglu, E. F. Plow, E. Gratton, and A. R. Horwitz, Integrin-associated complexes form hierarchically with variable stoichiometry in nascent adhesions, Curr. Biol. **24**, 1845 (2014).
[13] S. Yamashiro, D. M. Rutkowski, K. A. Lynch, Y. Liu, D. Vavylonis, and N. Watanabe, Force transmission by retrograde actin flow-induced dynamic molecular stretching of talin, Nat. Commun. **14**, 8468 (2023).
[14] K. Austen, P. Ringer, A. Mehlich, A. Chrostek-Grashoff, C. Kluger, C. Klingner, B. Sabass, R. Zent, M. Rief, and C. Grashoff, Extracellular rigidity sensing by talin isoform-specific mechanical linkages, Nat. Cell Biol. **17**, 1597 (2015).
[15] P. Ringer, A. Weißl, A.-L. Cost, A. Freikamp, B. Sabass, A. Mehlich, M. Tramier, M. Rief, and C. Grashoff, Multiplexing molecular tension sensors reveals piconewton force gradient across talin-1, Nat. Methods **14**, 1090 (2017).
[16] F. Kong, Z. Li, W. M. Parks, D. W. Dumbauld, A. J. García, A. P. Mould, M. J. Humphries, and C. Zhu, Cyclic mechanical reinforcement of integrin–ligand interactions, Mol. Cell **49**, 1060 (2013).
[17] J. Li, J. Yan, and T. A. Springer, Low-affinity integrin states have faster ligand-binding kinetics than the high-affinity state, eLife **10**, e73359 (2021).
[18] R. Oria, T. Wiegand, J. Escribano, A. Elosegui-Artola, J. J. Uriarte, C. Moreno-Pulido, I. Platzman, P. Delcanale, L. Albertazzi, D. Navajas, X. Trepat, J. M. García-Aznar, E. A. Cavalcanti-Adam, and P. Roca-Cusachs, Force loading explains spatial sensing of ligands by cells, Nature **552**, 219 (2017).
[19] M. H. Jo, P. Meneses, O. Yang, C. C. Carcamo, S. Pangeni, and T. Ha, Determination of single-molecule loading rate during mechanotransduction in cell adhesion, Science **383**, 1374 (2024).
[20] J. F. Beausang, H. W. Schroeder, 3rd, P. C. Nelson, and Y. E. Goldman, Twirling of actin by myosins II and V observed via polarized TIRF in a modified gliding assay, Biophys. J. **95**, 5820 (2008).
[21] L. Melli, N. Billington, S. A. Sun, J. E. Bird, A. Nagy, T. B. Friedman, Y. Takagi, and J. R. Sellers, Bipolar filaments of human nonmuscle myosin 2-A and 2-B have distinct motile and mechanical properties, eLife **7**, e32871 (2018).
[22] J. E. Molloy, J. E. Burns, J. Kendrick-Jones, R. T. Tregear, and D. C. S. White, Movement and force produced by a single myosin head, Nature **378**, 209 (1995).
[23] S. Hu, K. Dasbiswas, Z. Guo, Y.-H. Tee, V. Thiagarajan, P. Hersen, T.-L. Chew, S. A. Safran, R. Zaidel-Bar, and A. D. Bershadsky, Long-range self-organization of cytoskeletal myosin II filament stacks, Nat. Cell Biol. **19**, 133 (2017).
[24] N. Billington, A. Wang, J. Mao, R. S. Adelstein, and J. R. Sellers, Characterization of three full-length human nonmuscle myosin II paralogs, J. Biol. Chem. **288**, 33398 (2013).
[25] D. A. Calderwood, I. D. Campbell, and D. R. Critchley, Talins and kindlins: partners in integrin-mediated adhesion, Nat. Rev. Mol. Cell Biol. **14**, 503 (2013).
[26] R. S. Winardhi, Q. Tang, J. Chen, M. Yao, and J. Yan, Probing small molecule binding to unfolded polyprotein based on its elasticity and refolding, Biophys. J. **111**, 2349 (2016).
[27] X. Liu, Y. Wang, M. Yao, K. B. Baker, B. Klapholz, N. H. Brown, B. T. Goult, and J. Yan, The mechanical response of vinculin, bioRxiv (2023).
[28] S. Le, X. Hu, M. Yao, H. Chen, M. Yu, X. Xu, N. Nakazawa, F. M. Margadant, M. P. Sheetz, and J. Yan, Mechanotransmission and mechanosensing of human alpha-actinin 1, Cell Rep. **21**, 2714 (2017).
